# Functional redundancy between UTY and UTX in regulating the localization of transcription factors involved in pluripotency

**DOI:** 10.1101/2025.07.03.663017

**Authors:** Tomohiko Akiyama, Toshiya Nakahara, Saeko Sato, Kei-ichiro Ishiguro, Masashi Yukawa, Miu Yamamoto, Hidehisa Takahashi, Minoru S H Ko

## Abstract

The Y chromosome harbors few protein-coding genes, and their roles in early human development remain largely unclear. Here, we demonstrate that UTY, a Y-linked homolog of the histone demethylase UTX, plays a critical, demethylase-independent role in maintaining pluripotency in human embryonic stem cells (hESCs). Despite its low expression and weak enzymatic activity, UTY co-occupies active cis-regulatory elements with UTX and promotes the transcription of key pluripotency genes, including *NODAL* and *LEFTY*. Integrated genomic analyses of single and double knockout hESCs reveal that UTY and UTX function redundantly to maintain chromatin accessibility and ensure proper recruitment of core transcription factors such as OCT4 and SOX2. The combined loss of UTY and UTX disrupts transcription factor localization, induces widespread gene expression changes, and leads to a collapse of the pluripotent state, without detectable changes in H3K27 methylation. Instead, these defects are associated with impaired recruitment of ATP-dependent chromatin remodelers and reduced histone acetylation, suggesting a demethylase-independent mechanism. Our findings uncover an essential role for the evolutionarily conserved UTY in safeguarding enhancer function and transcription factor occupancy, highlighting Y-linked regulatory mechanisms that are critical for maintaining human pluripotency.

**Summary statement:** This study reveals how UTX and UTY cooperate to maintain stable gene regulation in human stem cells and guide proper developmental programs.

## Introduction

The human X and Y chromosomes have evolved from a pair of ancestral autosomes (Lahn and Page, 1999). While the X chromosome retains ∼98% of the original gene content, the Y chromosome has undergone substantial degeneration and preserved only ∼3% of ancestral genes (Bellott et al., 2010; Mueller et al., 2013; Skaletsky et al., 2003). Nevertheless, a select subset of Y-linked protein-coding genes has been evolutionarily conserved for at least 25 million years (Bellott et al., 2014; Hughes et al., 2012; Hughes et al., 2005). These genes encode proteins involved in key processes such as transcription, translation, and protein stability (Bellott et al., 2014; Hughes and Page, 2015), and many are broadly expressed across somatic tissues (Godfrey et al., 2020), suggesting roles beyond male reproduction. However, their precise biological functions remain poorly defined. Understanding their contributions to development and aging may offer valuable insights into the evolutionary significance of the Y chromosome’s gene repertoire.

Among these conserved Y-linked genes, UTY (Ubiquitously Transcribed Tetratricopeptide Repeat Gene on the Y chromosome) is of particular interest for its putative role in transcriptional regulation during early development, as supported by several lines of evidence.

UTY encodes a Jumonji C (JmjC) domain–containing protein, classified as a member of histone demethylase family (Klose et al., 2006). In vitro assays have indicated that UTY can demethylate trimethylated lysine 27 on histone H3 (H3K27), a repressive histone modification associated with gene silencing (Walport et al., 2014). However, UTY’s enzymatic activity is substantially reduced relative to its X-linked homolog UTX (KDM6A) and no demethylase activity has been detected in cultured cells (Agger et al., 2007; Hong et al., 2007). By contrast, UTX has been characterized as a bona fide H3K27 demethylase required for lineage specification and differentiation (Lee et al., 2012; Lei and Jiao, 2018; Mansour et al., 2012; Seenundun et al., 2010; Tang et al., 2020). Despite its limited catalytic activity, UTY is broadly expressed and has been conserved across mammals, raising the possibility that it fulfills a distinct, non-enzymatic regulatory role.

One potential role of UTY is to balance sex chromosome gene dosage between males and females. In preimplantation female embryos, both X chromosomes are transcriptionally active, resulting in elevated expression of X-linked genes relative to XY males. To restore dosage equilibrium, X-inactivation is initiated around the time of implantation, leading to the stable transcriptional silencing of one X chromosome. However, UTX escapes X-inactivation (Greenfield et al., 1998) and is expressed from both alleles in females throughout development, while males rely on the combined expression of UTX and UTY. Given UTY’s weak catalytic capacity, it remains unclear to what extent this co-expression provides functional compensation. Notably, Kabuki syndrome, a disorder caused by UTX haploinsufficiency and typified by growth delay and intellectual disability, is significantly more severe in males (UTX^–/Y) than in heterozygous females (UTX^+/–) (Banka et al., 2015; Lederer et al., 2014), indicating that UTY cannot fully substitute for UTX’s enzymatic function during late stages of development.

By contrast, UTY appears to play a more effective compensatory role during early embryogenesis. In mouse models, deletion of UTX leads to embryonic lethality in homozygous females by E11.5, whereas hemizygous UTX-null males—which retain UTY—survive to term, albeit at low frequency (∼20%) and with reduced body size (Lee et al., 2012; Shpargel et al., 2012; Thieme et al., 2013; Wang et al., 2012; Welstead et al., 2012). These sexually dimorphic phenotypes suggest that UTY and UTX act redundantly during early development, and that UTY’s presence is sufficient to support embryogenesis in the absence of UTX. Consistent with this, catalytic activity of UTX is dispensable for early lineage commitment, including in neural precursor and mesodermal differentiation (Shan et al., 2020; Wang et al., 2012). Together, these indicate that UTY plays a critical, demethylase-independent role in early embryonic development.

In this study, we revisit the role of UTY in male human embryonic stem (ES) cells, which harbor one allele each of UTX and UTY, and dissect their individual and combined contributions to pluripotency maintenance. By analyzing UTX and UTY single- and double-knockout lines, we demonstrate that UTY sustains enhancer activation, transcription factor occupancy, and an open chromatin state associated with pluripotency in the absence of UTX. These findings reveal a critical, non-catalytic role for UTY in preserving transcriptional networks during early human development, functionally paralleling UTX.

## Results

### UTY Is Weakly Expressed Relative to UTX in Human Pluripotent Stem Cells

Although UTY exhibits lower demethylase activity compared to UTX, the two proteins share highly similar structures (Fig. 1A). UTY is broadly expressed across various human cells and tissues, but its expression levels in pluripotent stem cells—particularly in comparison to UTX—have not been well characterized. To address this, we analyzed publicly available RNA-sequencing data from five pluripotent stem cell lines (male), including three ES cell lines (SEES3 (Nakatake et al., 2020), H1 (Wei et al., 2021), WIBR1 (Theunissen et al., 2016)) and two iPS cell lines (585A1, NPC1.1 (Theunissen et al., 2016)). We quantified the expression levels of UTX and UTY using normalized transcript counts (TPM: transcripts per kilobase million). UTY expression was consistently low but detectable (approximately 2–12 TPM) across all lines, while UTX was expressed at 3- to 4-fold higher levels (approximately 7–48 TPM) (Fig. 1B). These results indicate that although UTY is transcribed in human pluripotent stem cells, its expression is markedly lower than that of UTX.

**Figure 1.**
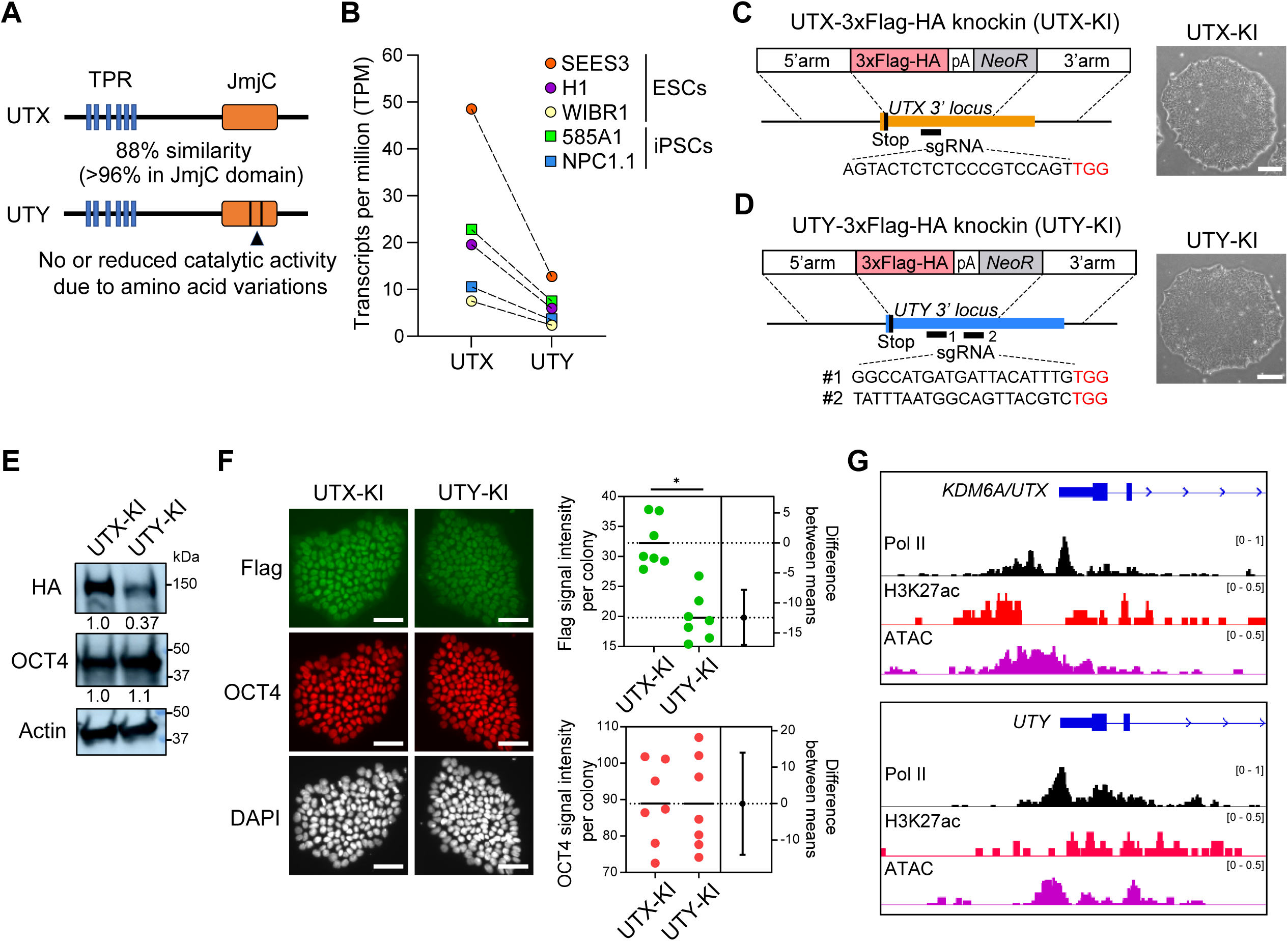
UTY is expressed at lower levels than UTX in human pluripotent stem cells. (A) Domain organization of UTX and UTY. TPR, tetratricopeptide repeat; JmjC, Jumonji C domain. (B) Comparison of RNA expression levels of UTX and UTY in human pluripotent stem cell lines. Transcripts per million (TPM) values are shown for ES cell lines (SEES3, H1, WIBR1) and iPS cell lines (585A1, NPC1.1). (C, D) CRISPR-Cas9 was used to generate UTX-3xFlag-HA and UTY-3xFlag-HA knock-in (UTX-KI and UTY-KI) human ES cells by inserting a 3xFlag-HA tag at the C-terminus of each gene. sgRNA target sequences are shown in black, and PAM sequences are shown in red. (E) Protein expression levels of UTX and UTY were assessed by immunoblotting using an HA antibody. ACTB antibodies were used as loading controls. The values indicate HA and OCT4 signal intensities normalized by ACTB signals. (F) Immunostaining analysis using a Flag antibody in UTX-KI and UTY-KI cells. The signal intensities of ES colonies (*N*L=L7) were quantified. \**P*L<L0.01 by one-sided Student’s t-test. The OCT4 staining intensities were used as a control. Nuclei were counterstained with DAPI. (G) Profiles of Pol II, H3K27ac, and ATAC-seq signals at the promoter regions of the UTX and UTY genes.

To directly compare UTX and UTY protein levels under identical conditions, we generated human ES cell lines in which a 3×Flag-HA epitope tag was inserted at the C-terminus of the endogenous *UTX* or *UTY* loci using CRISPR-Cas9 genome editing (Fig. 1C,D, and Fig. S1). These knock-in cell lines, referred to as UTX-KI and UTY-KI, were derived from the SEES3 cell line, which exhibits the highest transcript levels among those examined. Immunoblotting with an HA antibody in these cell lines revealed that UTY protein levels were significantly lower than those of UTX (FC = 0.37) (Fig. 1E). Similarly, immunostaining with a Flag antibody confirmed the reduced levels of UTY compared to UTX (FC = 0.66) (Fig. 1F). Expression of the pluripotency marker OCT4 was comparable between UTX-KI and UTY-KI cells.

These results demonstrate that although UTY is expressed in human ES cells, its protein levels are substantially lower than those of UTX. We hypothesized that this difference might arise from transcriptional regulation. However, comparative analysis of RNA polymerase II occupancy, H3K27 acetylation, and chromatin accessibility at the *UTX* and *UTY* promoters showed no significant differences (Fig. 1G). This suggests that the reduced expression of UTY is unlikely to be due to differences in transcriptional activation, and may instead result from lower RNA stability, shorter protein half-life, or other epigenetic mechanisms.

### UTY is Co-localized with UTX at Promoters and Enhancers

To investigate the potential role of the weakly expressed UTY alongside UTX, we examined the genome-wide binding profiles of both proteins. We performed chromatin immunoprecipitation followed by sequencing (ChIP-seq) using an anti-Flag antibody in UTX-KI and UTY-KI cell lines. During optimization of the experimental conditions, we found that formaldehyde crosslinking alone was insufficient to capture ChIP-seq peaks, likely because UTX and UTY do not bind DNA directly. To overcome this, we used a dual crosslinking method with disuccinimidyl glutarate prior to formaldehyde fixation, which enhances the crosslinking of protein-DNA complexes (Tian et al., 2012). This approach improved data quality, yielding distinct and reproducible peaks for both proteins (Fig. 2A). Consistent with their expression levels, we identified ∼37,000 UTX peaks and ∼10,000 UTY peaks across biological triplicates (Fig. 2B). Despite this difference in peak number, both proteins showed similar genomic distributions: they were enriched at core promoters (0–0.5 kb), proximal regulatory regions (0.5–1 kb), and distal regulatory elements within ∼50 kb of transcription start sites (TSS), likely corresponding to enhancers (Fig. 2B).

**Figure 2.**
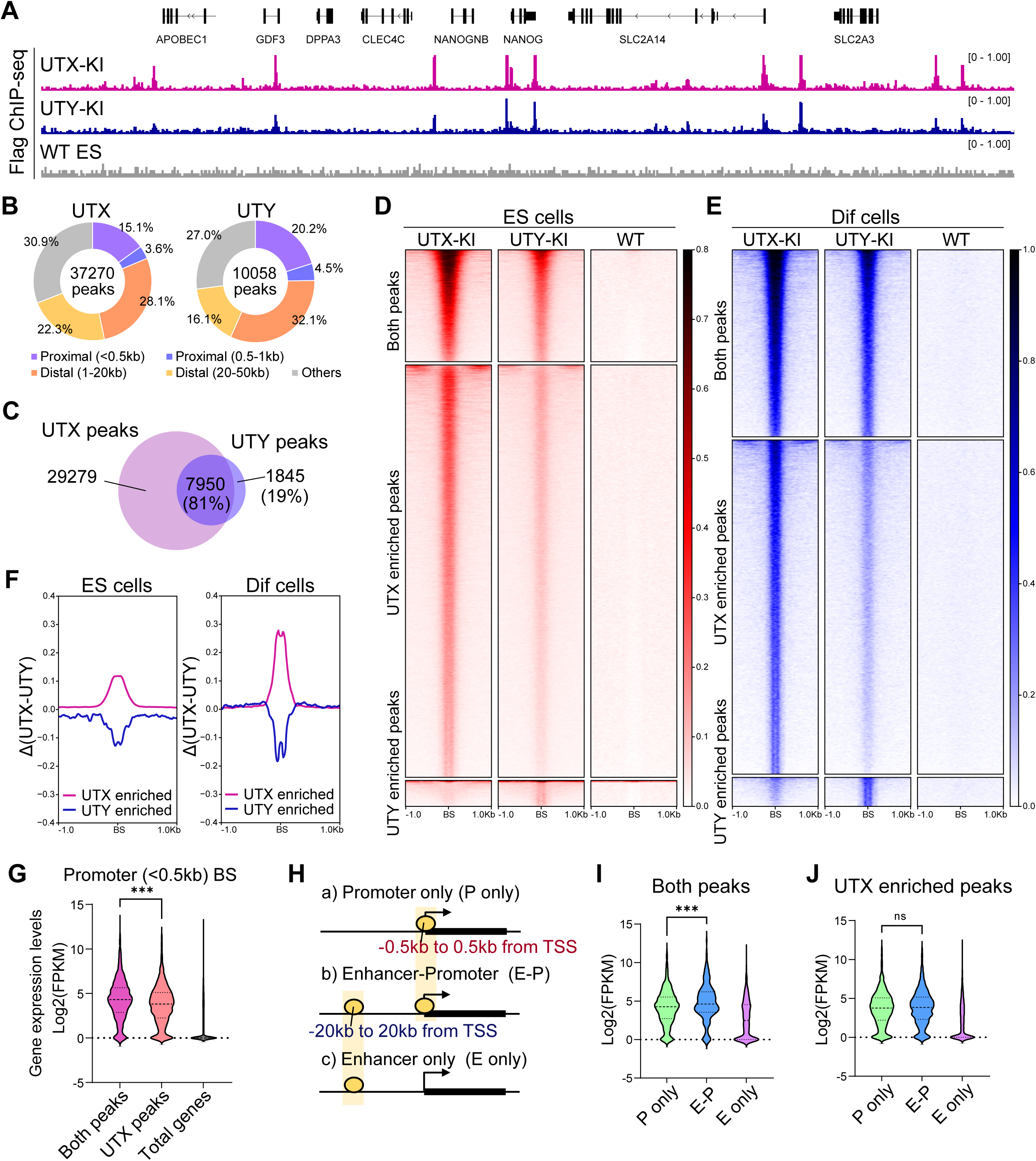
UTY is co-localized with UTX at promoters and enhancers in active genes. (A) ChIP-seq tracks representing UTX and UTY localization patterns in ES cells. ChIP-seq analysis was performed using a Flag antibody in UTX-KI and UTY cells. Wild-type ES cells were used as a negative control. (B) Number of the UTX and UTY ChIP-seq peaks and their distribution relative to the TSS. (C) Overlapping of the UTX and UTY ChIP-seq peaks. (D) Flag ChIP-seq signal intensities of UTX-KI and UTY-KI cells are shown within a 2 kb region centered on the sites of both peaks, UTX-enriched peaks, and UTY-enriched peaks in ES cells. Flag ChIP-seq signal intensities of wild-type ES cells were used as a negative control. (E) ChIP-seq signal intensities of UTX-KI and UTY-KI within a 2 kb region centered on the sites of both peaks, UTX-enriched peaks, and UTY-enriched peaks in differentiated cells. Wild-type ES cells were used as a negative control. (F) Differences of UTX and UTY ChIP-seq signal intensities at UTX-enriched sites and UTY-enriched sites in ES cells and differentiated cells. (G) Gene expression levels of genes with both peaks or UTX-enriched peaks at promoter regions. Total genes are used as a control. \**P*L<L0.01 by one-sided Student’s t-test. (H) Localization patterns related to gene regulation: a) promoter only, b) enhancer and promoter, c) enhancer only. (I) Gene expression levels of genes with both peaks at the three localization patterns. \**P*L<L0.01 by one-sided Student’s t-test. (J) Gene expression levels of genes with UTX peaks at the three localization patterns. ‘ns’ indicates not significant.

UTX and UTY binding regions substantially overlapped, with ∼80% of UTY peaks co-localizing with UTX peaks (Fig. 2C). We define these co-localized regions as UTX/UTY co-bound regions (“Both-enriched peaks,” hereafter referred to as “Both peaks”). Heatmap analysis of ChIP-seq signals showed that these regions exhibit relatively high enrichment of both UTX and UTY across all binding sites (Fig. 2D). Outside of these co-bound regions, comparative analysis revealed a greater number of UTX-specific peaks (Fig. 2C). However, UTY signals were still detectable at most of these sites, although they did not exceed the peak-calling threshold (Fig. 2D); we designated these as “UTX-enriched sites.” This indicates that most UTX-bound regions are also occupied by UTY at lower levels. In contrast, only a small subset of UTY peaks showed minimal or no UTX signal and were defined as “UTY-enriched sites” (Fig. 2C,D). Together, these results suggest that UTY primarily functions in concert with UTX in human ES cells.

The co-localization of UTX and UTY was also observed in differentiated cells (endoderm cells derived from ES cells after 5 days of induction) (Fig. 2E). Most UTY signals overlapped with UTX, and low levels of UTY were detected at many UTX-enriched sites, whereas UTY-enriched sites were relatively few. These results indicate that UTX and UTY generally co-occupy similar genomic regions even in differentiated cells. Although UTY-enriched sites were detected, their minimal overlap between undifferentiated and differentiated cells indicates that these sites are not fixed across cell states. Notably, the difference in binding levels between UTX and UTY—denoted as Δ(UTX−UTY)—was more pronounced in differentiated cells than in undifferentiated ES cells (Fig. 2F), suggesting that their functional divergence may emerge during differentiation.

To investigate the functional relevance of UTX and UTY co-occupancy in human ES cells, we analyzed gene expression associated with promoter-bound peaks. Most genes with promoter binding were actively transcribed, and those bound by both UTX and UTY exhibited higher transcriptional activity than those bound by UTX alone (Fig. 2G). These results suggest that robust binding and co-occupancy of UTX and UTY is associated with enhanced transcriptional output. We performed similar analyses for UTY-enriched sites, but their low number and weak signal intensity limited the ability to draw definitive conclusions.

We further examined gene expression associated with distal UTX/UTY binding sites. Genes were grouped into three categories: those with binding exclusively at promoters, exclusively at distal regions (putative enhancers), and at both promoters and enhancers (Fig. 2H). Genes with both promoter and enhancer binding showed the highest expression levels (Fig. 2I). In contrast, genes with binding sites restricted to enhancers displayed both active and inactive states. A similar trend was observed for genes associated with UTX-enriched sites, although their expression levels were generally lower than those linked to co-bound regions (Fig. 2I,J). These results suggest that UTX–UTY co-occupancy promotes more robust and defined transcriptional regulation.

In differentiated cells, UTX and UTY were also detected at both promoters and distal regions (Fig. S2A,B). These binding patterns were distinct from those of JMJD3, another H3K27 demethylase that functions during ES cell differentiation (Akiyama et al., 2016; Burgold et al., 2008; Kartikasari et al., 2013). JMJD3 showed broad distribution from promoters to gene bodies, consistent with its catalytic role in H3K27 demethylation (Fig. S2C,D), whereas UTX and UTY exhibited more discrete binding patterns. This aligns with previous findings showing that forced expression of JMJD3 alters H3K27 methylation in human ES cells, while UTX overexpression does not (Akiyama et al., 2016). These results suggest that the genomic localization of UTX and UTY reflects non-catalytic functions rather than enzymatic activity. Interestingly, UTX/UTY co-bound peak sites showed a higher degree of overlap between ES and differentiated cells compared with UTX-only peaks (13.4% vs. 6.6% of ES peaks, respectively), indicating that shared sites are more stably maintained during early differentiation, whereas UTX-only binding is more dynamically regulated. Gene Ontology analysis further revealed that genes associated with UTX/UTY co-bound peaks are enriched for transcriptional and RNA regulatory processes, whereas UTX-enriched peaks are preferentially linked to neurodevelopment-related genes, consistent with lineage priming in human ES cells (Fig. S3).

### Identification of Genes Cooperatively Regulated by UTX and UTY

To further assess the functional redundancy of UTX and UTY, we generated human ES cell lines with single (UTX-KO and UTY-KO) and double (DKO) deletions using CRISPR-Cas9. Gene-specific deletions and homozygous mutations were confirmed in single and double knockout lines, respectively (Fig. S4). In single knockout cells, a modest increase in the protein level of the remaining paralog was observed, without a corresponding increase in mRNA levels.

We performed RNA sequencing (RNA-seq) on these cell lines to evaluate transcriptomic changes. We first examined expression changes across all genes and found that single KO of either UTX or UTY resulted in minimal global changes, whereas DKO cells exhibited pronounced transcriptional alterations (Fig. S5A). We then focused on genes located within 20 kb of UTX and/or UTY binding sites to assess more direct regulatory effects. Genes were grouped according to their binding patterns: UTX-enriched, UTY-enriched, or co-occupied (“both-enriched”). Under single KO conditions, only a small subset of genes was differentially expressed, whereas DKO cells showed a markedly larger number of changes (Fig. 3A,B), with downregulation being more frequent—supporting a role for UTX and UTY in transcriptional activation.

**Figure 3.**
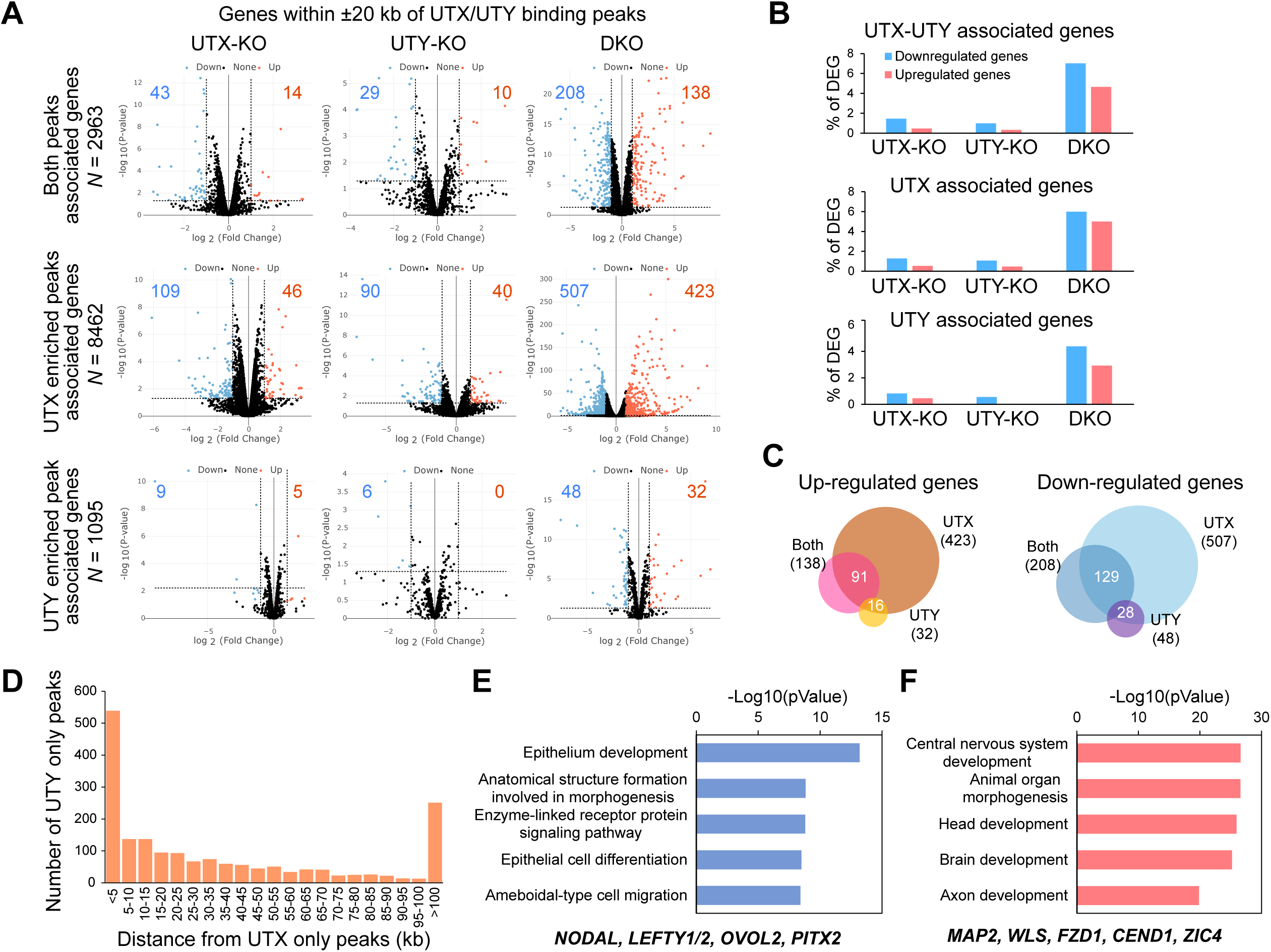
Genes expression changes in UTX-KO, UTY-KO, and DKO cells. (A) Gene expression changes in UTX-KO, UTY-KO, and DKO cells are shown with volcano plots. Blue plots indicate downregulated genes, and red plots indicate upregulated genes. Th graphs are shown separately for genes associated with both peaks, UTX-enriched peaks, and UTY-enriched peaks (genes within 20 kb of the peaks). (B) The percentage of differentially expressed genes (DEGs) are summarized. (C) Overlapping of DEGs associated with both, UTX-enriched, and UTY-enriched peaks. (D) The relationship between UTX peaks and UTY peaks. A graph showing the number of UTY peaks located at the indicated distances from UTX peaks. (E) Characterization of downregulated genes in DKO cells. Example genes are shown. (F) Characterization of upregulated genes in DKO cells. Example genes are shown.

Furthermore, many genes with altered expression in DKO cells were shared among the three binding groups (Fig. 3C), suggesting cooperative regulation by UTX and UTY. Supporting this, UTX- and UTY-enriched sites were frequently located in close proximity, with a large proportion within 5 kb of each other (Fig. 3D). These findings indicate that UTX and UTY often co-occupy the same or adjacent genomic regions, enabling coordinated transcriptional regulation.

Gene ontology analysis revealed that genes downregulated in DKO cells were significantly enriched for biological processes reflecting the epithelial-like characteristics of pluripotent cells and featured key regulators such as *NODAL* and *LEFTY* (Fig. 3E). Conversely, genes upregulated in DKO cells were associated with neural differentiation pathways, including *MAP2* and *WLS* (Fig. 3F).

Collectively, these results indicate that UTX and UTY do not function independently in human ES cells, but instead act redundantly to sustain the expression of pluripotency-associated genes and to prevent premature activation of differentiation programs.

To further examine whether UTY compensates for UTX in a sex-dependent manner, we generated UTX-KO female human ES cells and performed RNA-seq analysis. In contrast to the minimal transcriptomic changes observed in male UTX-KO cells, UTX loss in female cells induced gene expression changes that were highly similar to those observed in DKO male cells (Fig. S5B). These findings suggest that UTY in male ES cells can compensate for UTX loss, functionally replacing the extra UTX dosage present in female cells.

### UTX and UTY Regulate the Binding Abilities of OCT4 and SOX2 at Enhancers

The co-occupancy of UTX and UTY at active enhancers prompted us to examine whether their presence influences the binding of OCT4 and SOX2, two core transcription factors essential for maintaining pluripotency. Consistent with this idea, UTX- and UTY-bound regions were highly enriched for OCT4 and SOX2 binding motifs (Fig. 4A). ChIP-seq analysis further showed enrichment of OCT4 and SOX2 signals at regions co-bound by UTX and UTY (Both sites), as well as at UTX-enriched sites (Fig. 4B). Notably, a subset of UTX/UTY-bound regions which lacked OCT4 and SOX2 occupancy was predominantly associated with promoters (Fig. 4C), suggesting that UTX and UTY bind both enhancers and promoters and may link enhancer-driven transcription factor activity to promoter regulation. When comparing WT and DKO cells, global OCT4 and SOX2 occupancy patterns appeared largely preserved, with only a modest overall decrease (Fig. 4B). We therefore focused our analysis on OCT4 and SOX2 binding at enhancers. At enhancers co-bound by UTX and UTY within 20 kb of the nearest TSS, OCT4 binding was reduced in DKO cells compared with WT (Fig. 4D). This reduction was more pronounced at enhancers associated with genes downregulated in DKO cells (Fig. 4E,F). A similar decrease was observed for SOX2 binding (Fig. 4G–I), particularly at enhancers linked to key pluripotency genes such as *LEFTY* and *NODAL* (Fig. 4J,K). Additionally, genes downregulated in DKO cells showed significantly reduced RNA polymerase II loading at their promoters (Fig. S6). Notably, these changes occurred with minimal alterations in H3K27me3 levels (Fig. S7). These data indicate that loss of UTX and UTY compromises OCT4/SOX2 occupancy and promoter Pol II loading without major changes in H3K27me3.

**Figure 4.**
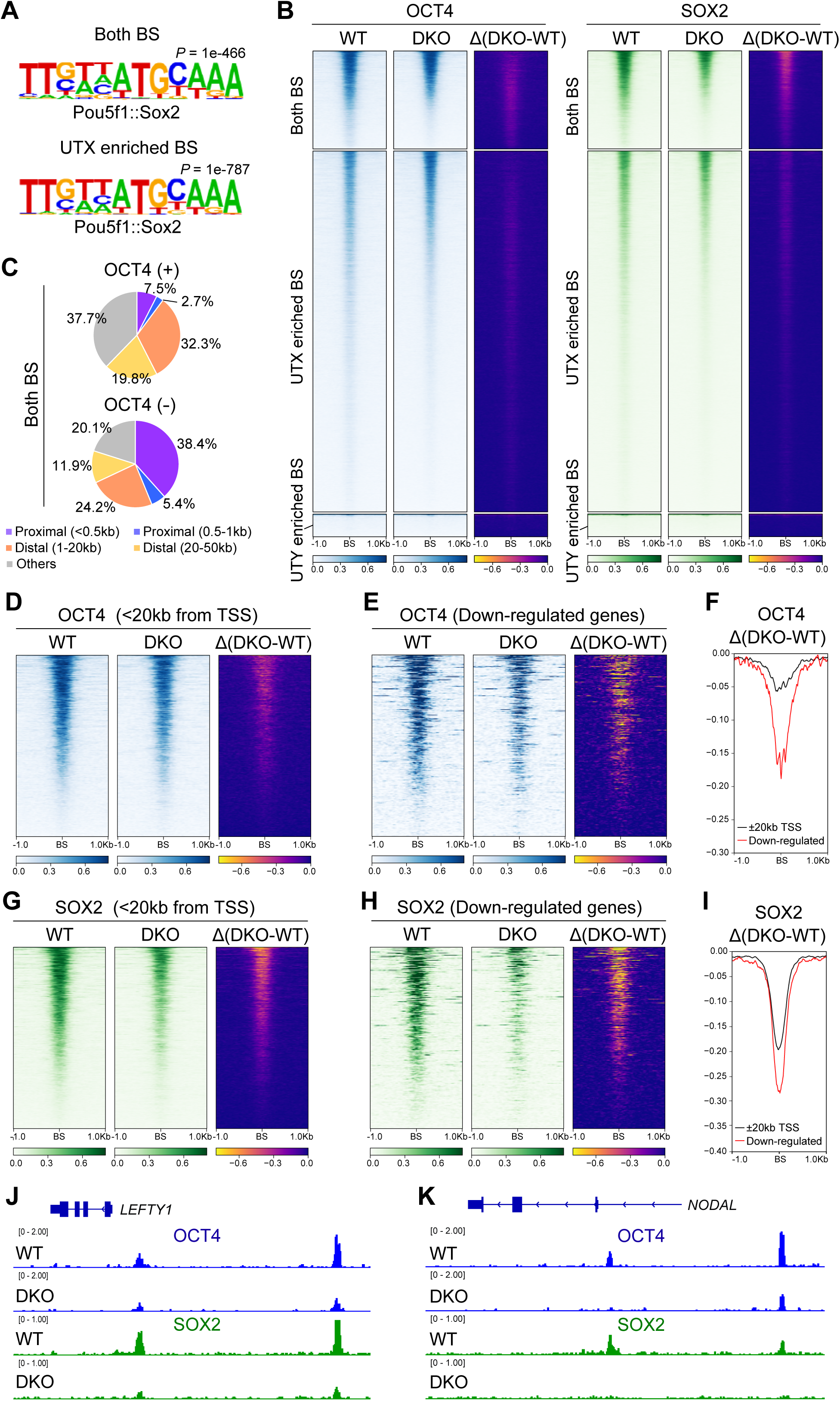
UTX/UTY DKO impairs the binding of OCT4 and SOX2 at active enhancers. (A) Enriched sequence motifs at both UTX/UTY binding sites (BS) and UTX enriched BS. (B) ChIP-seq signal intensities of OCT4 and SOX2 centered (±1 kb) on UTX/UTY (Both), UTX-enriched, and UTY-enriched BS in WT and DKO cells. Δ(DKO−WT) represents the difference in normalized ChIP–seq signal intensity. (C) Genomic distribution of Both BS relative to the TSS with or without OCT4 co-occupancy. (D) OCT4 ChIP-seq signals in WT and DKO cells centered on Both BS located within 20 kb of TSS. Signal differences between WT and DKO also shown. (E) Same as (D), but restricted to BS near genes downregulated in DKO cells. (F) Comparison of OCT4Δ(DKO–WT) signals at Both BS within 20 kb of the TSS of all genes versus downregulated genes. (G) SOX2 ChIP-seq signals in WT and DKO cells centered on Both BS within 20 kb of TSS, with differences shown. (H) Same as (G), but focused on sites near downregulated genes. (I) Comparison of SOX2Δ(DKO–WT) signals within 20 kb of the TSS of all genes versus downregulated genes. (J, K) Reduced OCT4 and SOX2 binding at the *LEFTY1* and *NODAL* loci in DKO cells.

### Depletion of UTX and UTY Results in the Relocation of OCT4 and SOX2

Given that deletion of UTX and UTY reduces OCT4 and SOX2 binding at downregulated genes, we expected the resulting transcriptional changes to resemble those caused by OCT4 depletion. However, comparison of RNA-seq data from DKO cells and OCT4 siRNA-treated cells (Akiyama et al., 2018) revealed minimal overlap in differentially expressed genes (Fig. 5A), suggesting that UTX/UTY loss does not simply reduce OCT4 levels. Indeed, OCT4 expression remained unchanged in UTX/UTY-depleted cells (Fig. 5B).

**Figure 5.**
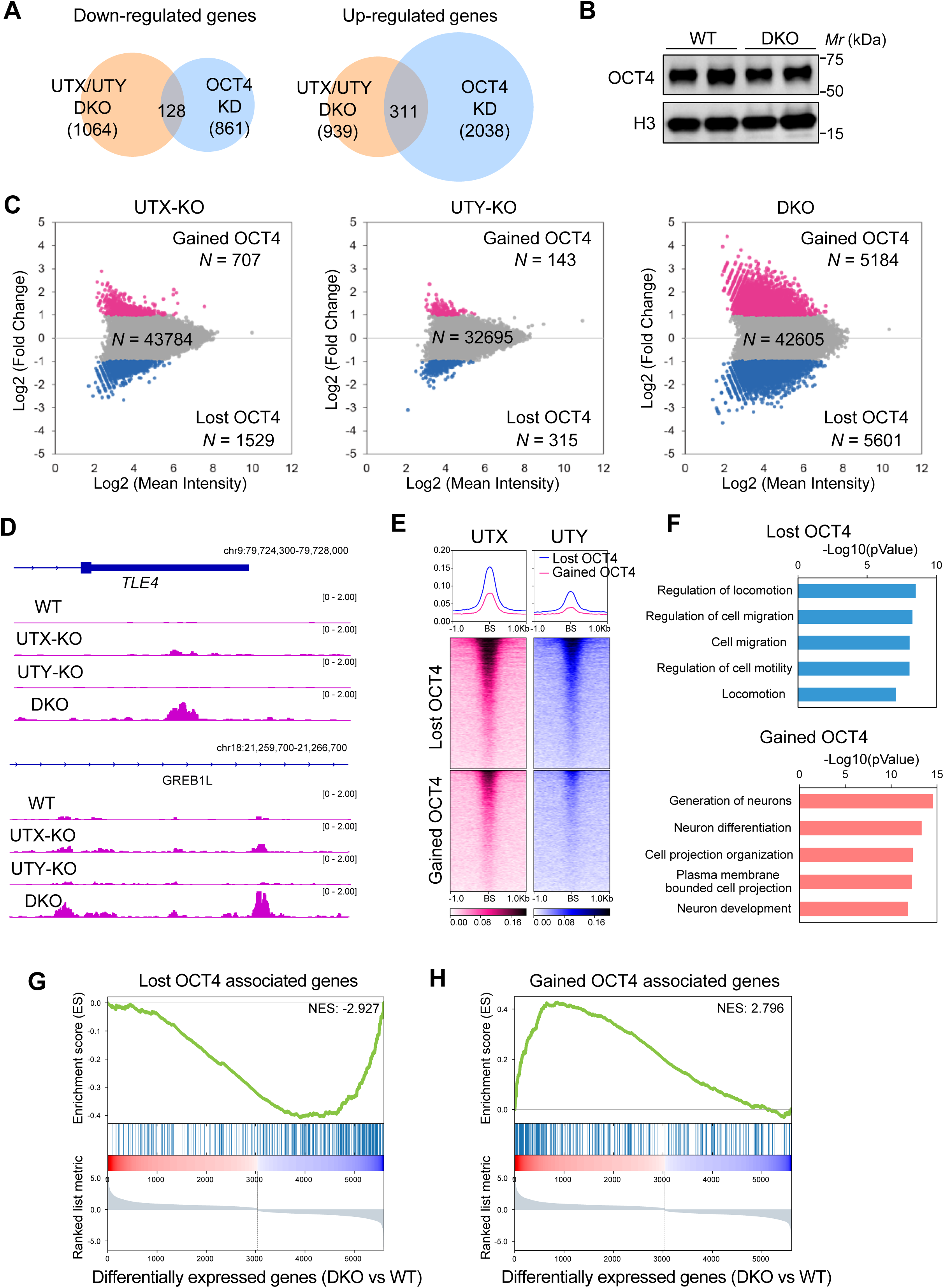
UTX/UTY DKO results in the relocation of OCT4. (A) Comparison of differentially expressed genes between UTX/UTY DKO cells and OCT4 KD (knockdown) cells. (B) Immunoblotting analysis showing that OCT4 levels remain unchanged between WT and DKO cells. Histone H3 is used as a loading control. (C) Differences in OCT4 distribution patterns in UTX-KO, UTY-KO, and DKO cells compared to WT cells. Increased ChIP-seq peaks, indicating gained OCT4 binding, are shown in pink, while decreased ChIP-seq peaks, indicating lost OCT4 binding, are shown in blue. (D) Representative gene loci showing increased OCT4 occupancy in DKO cells compared with WT cells. (E) UTX and UTY enrichment at sites with lost and gained OCT4 BS. (F) Characterization of genes associated with lost and gained OCT4 BS (<20kb from TSS). (G,H) The relationship between genes associated with altered OCT4 levels and genes whose expression is changed by DKO. Lost OCT4 site-associated genes are enriched in genes downregulated in DKO cells (G), whereas gained OCT4 site-associated genes are enriched in genes upregulated in DKO cells (H).

We therefore investigated whether UTX/UTY loss alters OCT4 genomic binding. Analysis of OCT4 ChIP-seq data using a MA-plot tool (Shao et al., 2012) revealed that although the total number of OCT4 peaks remained largely unchanged, both peak intensities and their genomic locations were dynamically altered in DKO cells (Fig. 5C). Peaks with >2-fold change were categorized as “lost” or “gained” peaks. OCT4 was redistributed to regions normally unbound in wild-type ES cells—including gene loci such as *TLE4* and *GREB1L*, which are upregulated in DKO cells (Fig. 5C,D). Although some redistribution occurred in single KO cells, it was much more pronounced in DKO cells. A similar pattern was observed for SOX2 (Fig. S8). OCT4 loss preferentially occurred at sites enriched for UTX and UTY binding, whereas gained OCT4 sites showed relatively low UTX/UTY occupancy (Fig. 5E). Furthermore, we found that gained OCT4 binding events preferentially occurred near neuronal gene loci (Fig. 5F), consistent with the activation of neuron-associated transcriptional programs observed in DKO cells. Together, these results indicate that UTX and UTY contribute to maintaining proper genome-wide localization of core transcription factors and are critical for sustaining pluripotency.

To investigate how these altered binding dynamics relate to transcriptional changes, we compared the locations of lost and gained OCT4 peaks with nearby (<20 kb) gene expression changes. Lost OCT4 peaks were frequently located near genes downregulated in DKO cells (Fig. 5G), while gained peaks were associated with upregulated genes (Fig. 5H). These results suggest that UTX and UTY depletion leads to a redistribution of OCT4 binding that directly influences the expression of its target genes.

### UTX and UTY Regulate the Localization of Chromatin Remodeling Factors

To explore the mechanisms underlying OCT4 and SOX2 relocation, we examined the distribution of epigenetic and chromatin remodeling factors in DKO cells, focusing on MLL4 (a histone methyltransferase and core component of the COMPASS complex), BRG1 (the catalytic subunit of the SWI/SNF complex), and CHD7 (an ATP-dependent chromatin remodeler). In DKO cells, the occupancy of these factors was globally reduced across UTX/UTY-bound regions (Fig. S9A). Notably, their enrichment was significantly decreased at sites where OCT4 binding was lost (Fig. 6A,C), whereas it was increased at sites where OCT4 binding was gained (Fig. 6B,D). A similar pattern was observed at SOX2 relocation sites (Fig. S9B). Together, these results suggest that UTX and UTY are required to maintain proper localization of chromatin regulatory factors at enhancers.

**Figure 6.**
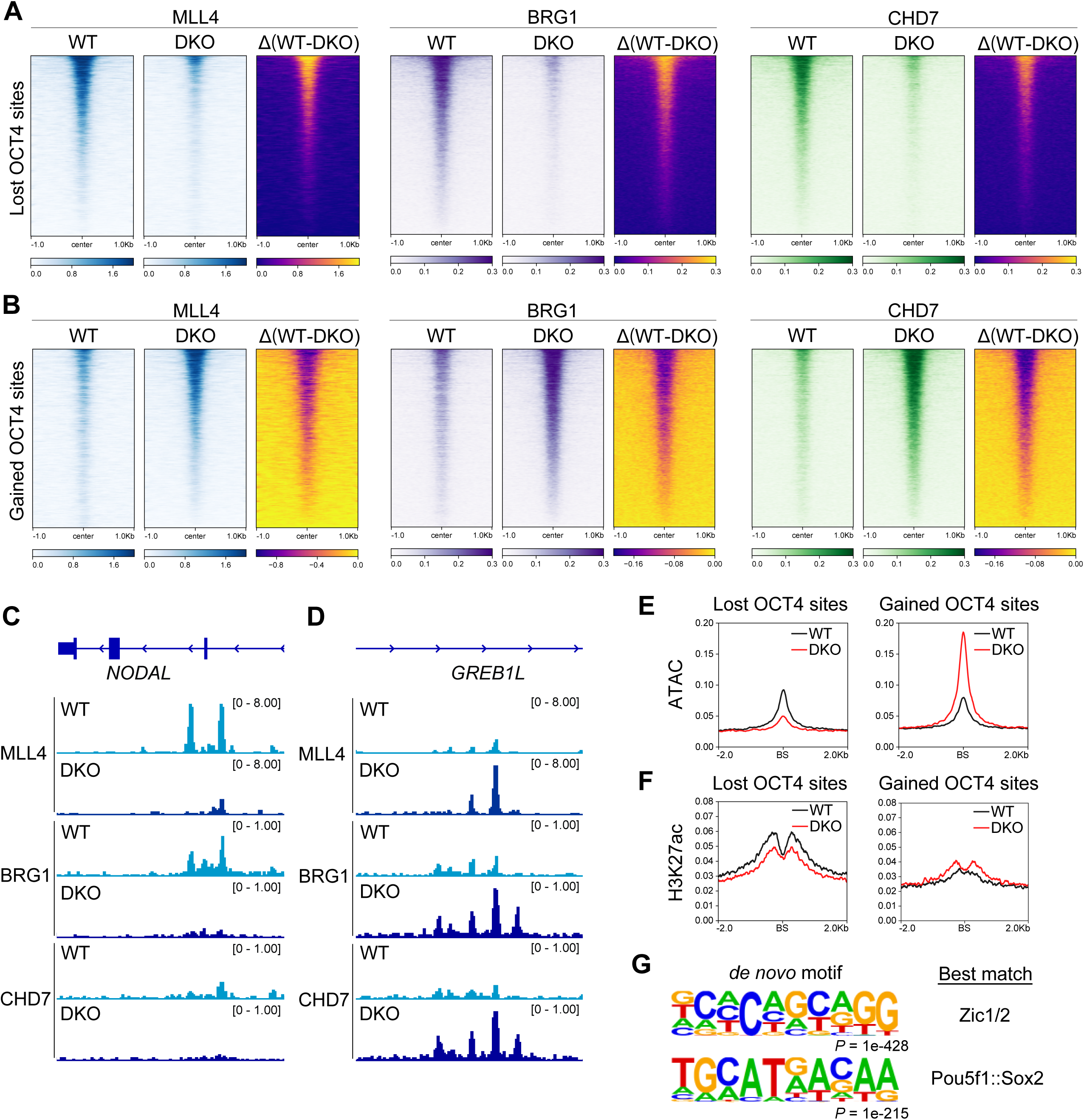
UTX and UTY are required for proper occupancy of MLL4, BRG1, and CHD7. (A) Comparison of ChIP-seq signal intensities of the indicated factors between WT and DKO cells centered on OCT4 lost sites. (B) Comparison of ChIP-seq signal intensities of the indicated factors between WT and DKO cells centered on OCT4 gained sites. (C) Example of a region showing reduced binding of the indicated factors. (D) Example of a region showing increased binding of the indicated factors. (E) ATAC-seq signals at OCT4 relocation sites in WT and DKO cells. (F) H3K27ac ChIP-seq signals at OCT4 relocation sites in WT and DKO cells. (G) Enriched sequence motifs at OCT4 gained sites.

Consistent with these findings, ATAC-seq and H3K27ac ChIP-seq analyses revealed parallel changes in chromatin accessibility and active enhancer marks (Fig. 6E,F), indicating that alterations in chromatin state likely contribute to transcription factor redistribution. Importantly, H3K27me3 levels remained largely unchanged at both lost and gained OCT4 sites (Fig. S10), supporting the conclusion that UTX and UTY function largely independently of their histone H3K27 demethylase activity.

The aberrant OCT4 binding observed in UTX- and UTY-deficient cells is likely influenced by the intrinsic properties of the affected genomic regions. Motif analysis of these aberrant OCT4-binding sites revealed enrichment of ZIC1/2 and POU5F1 (OCT4)/SOX2 motifs (Fig. 6G), suggesting these regions are inherently more accessible to transcription factors. This accessibility may be further increased by chromatin opening caused by UTX and UTY loss, thereby facilitating OCT4 binding.

Together, depletion of UTX and UTY leads to a detrimental loss of enhancer activity and promotes the formation of aberrant active enhancers by impairing the recruitment of chromatin remodeling factors.

### UTX and UTY Are Required for Maintenance of Human Pluripotency

Compared with WT and single KO cells, DKO cells exhibited subtle morphological changes and reduced alkaline phosphatase staining, a hallmark of undifferentiated ES cells (Fig. 7A). Immunostaining further revealed aberrant expression of the neuronal marker NESTIN despite largely preserved OCT4 expression, indicating a compromised pluripotent state with features of lineage priming (Fig. 7B). Under standard ES cell culture conditions, DKO cells remained proliferative, with overall growth comparable to that of WT and single KO cells (Fig. 7C). Upon in vitro differentiation, DKO cells upregulated representative markers of all three germ layers, indicating preserved responsiveness to differentiation cues (Fig. S11A). However, transcriptome analyses revealed distinct global gene expression patterns compared with WT and single KO cells, consistent with altered lineage specification programs (Fig. S11B).

**Figure 7.**
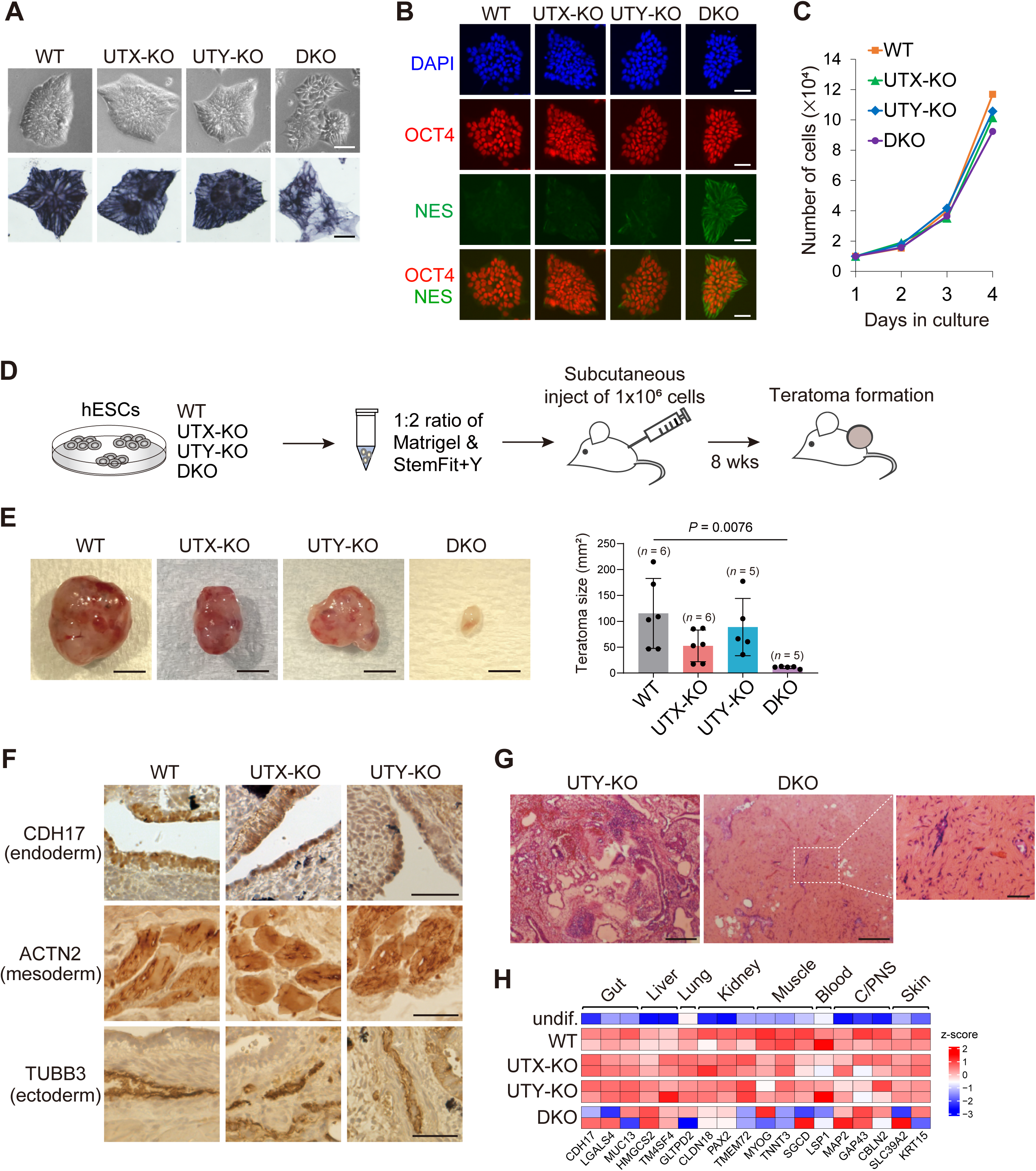
UTX and UTY are essential for sustaining pluripotency. (A) Representative bright-field images and alkaline phosphatase staining. Scale bar, 50 µm. (B) Immunostaining analysis of OCT4 and NESTIN. Scale bar, 50 µm. (C) Cell counts over 4 days. (D) Schematic representation of the teratoma formation assay to assess pluripotency in vivo. (E) Representative images of teratomas derived from WT, UTX-KO, UTY-KO, and DKO ES cells. A graph showing the measurement of the teratoma sizes. (F) Immunohistochemical staining with the indicated antibodies on the teratoma sections showing the differentiation into the three germ layers in WT and single KO conditions. (G) Hematoxylin and eosin staining showing that tissue-like structures were rarely observed in the DKO-derived masses. (H) Expression of marker genes for each tissue type in the teratomas.

In line with these findings, teratoma formation assays further supported impaired pluripotency upon combined loss of UTX and UTY (Fig. 7D). Eight weeks after injection into recipient mice, we analyzed the resulting teratomas/masses for morphology, tissue organization, and gene expression. While single KO cells developed teratomas, DKO cells failed to develop recognizable teratomas and only formed small, pale cell masses (Fig. 7E). These results suggest that UTX and UTY are important for maintaining human pluripotency in vivo. Notably, teratomas derived from UTX-KO and UTY-KO cells tended to be smaller than those from WT cells, with the size reduction more pronounced in UTX-KO teratomas, suggesting that differences in demethylase activity between UTX and UTY may begin to exert an influence during teratoma formation.

Histological analysis of the tissue sections revealed that single KO cells generated diverse tissues, including intestinal epithelium, muscle, and neural tubes, whereas DKO cells predominantly differentiated into stromal-like cells (Fig. 7F,G). Further analysis of gene expression related to the three germ layers showed that many genes failed to be upregulated in DKO cells, indicating a marked loss of pluripotency (Fig. 7H). These results suggest that enhancer regulation by UTX and UTY is essential for maintaining pluripotency and supporting human development.

## Discussion

In this study, we identified the genome-wide binding sites and downstream targets of both UTY and UTX, uncovering their redundant roles in transcriptional regulation in human ES cells. Both proteins regulate the recruitment of chromatin remodeling complexes to enhancers, thereby facilitating proper transcription factor occupancy and maintaining the pluripotent state (Fig. 8).

**Figure 8.**
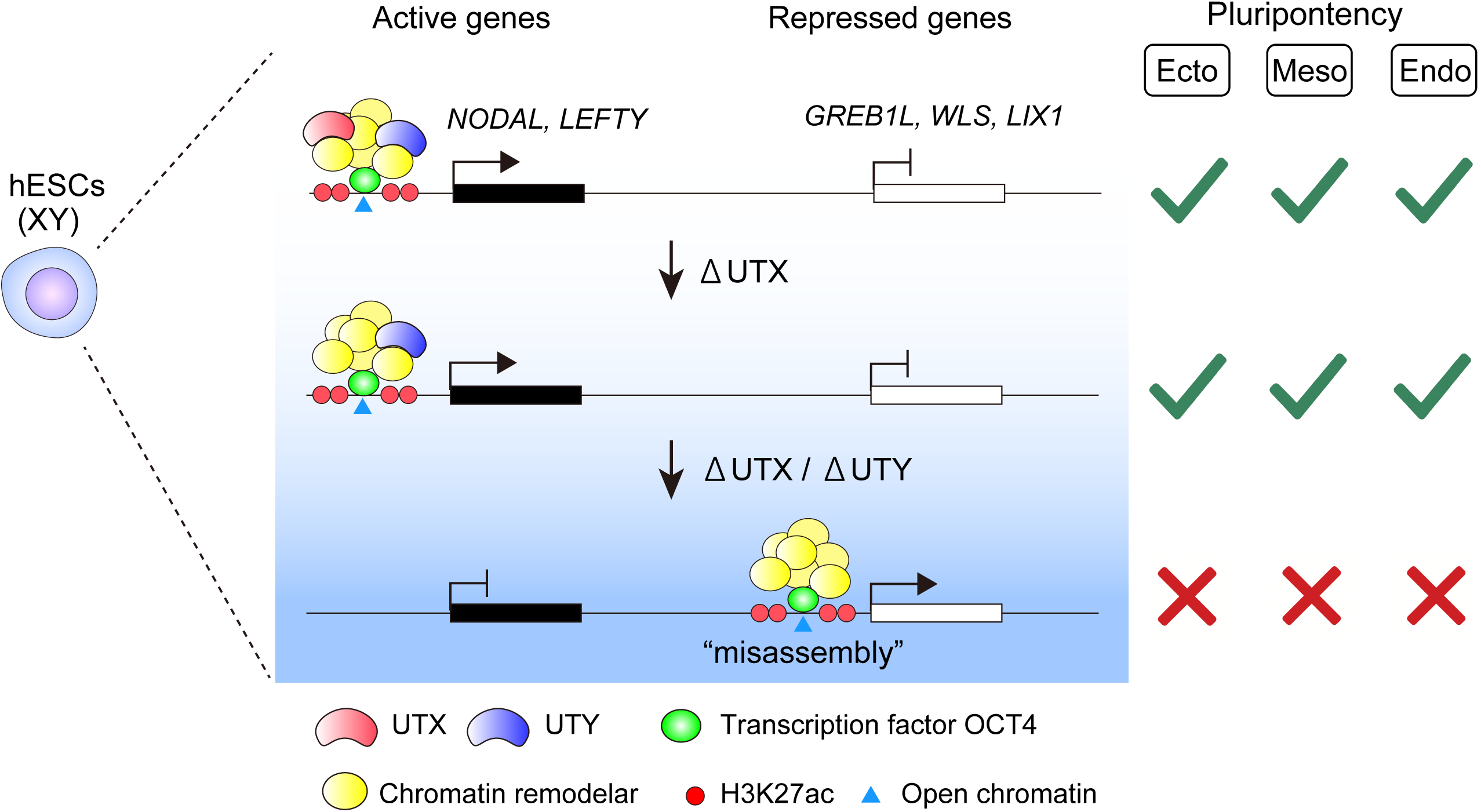
Model for UTY/UTX-dependent maintenance of transcription factor occupancy and pluripotency. Schematic representation of the model illustrating UTY’s role in facilitating transcription factor occupancy and maintaining pluripotency. In the absence of UTX, UTY is sufficient to preserve chromatin structure and maintain proper transcriptional regulation. However, when both UTX and UTY are absent, transcription factors and chromatin-associated complexes mislocalize—likely binding to alternative motifs—resulting in regulatory misassembly, ectopic activation of silent genes, and loss of pluripotency.

UTY, originally identified as a male-specific antigen (Greenfield et al., 1996), has long lacked a clearly defined function. By contrast, UTX is a well-characterized histone demethylase known to play essential roles in transcriptional activation. This functional distinction is reflected in their disease associations: UTX mutations are linked to disorders such as Kabuki syndrome (Banka et al., 2015; Lederer et al., 2012; Paderova et al., 2016) and acute lymphoblastic leukemia (Benyoucef et al., 2016; Mar et al., 2012; Ntziachristos et al., 2014; Van der Meulen et al., 2015), whereas comparable links for UTY are lacking. Despite these differences, our results demonstrate that UTX and UTY perform overlapping roles in chromatin regulation in human ES cells. In the context of pluripotency maintenance, the major transcriptional and chromatin phenotypes observed in DKO cells occur without substantial changes in H3K27me3 levels, suggesting that demethylase activity is largely dispensable for these functions.

Our findings show that UTX and UTY serve as linchpins in stabilizing the binding of transcription factors and chromatin remodeling factors. In the absence of both UTX and UTY, these factors become mislocalized, resulting in impaired enhancer activation and disruption of the gene regulatory network. Crucially, these functions are largely independent of changes in H3K27me3 levels. Rather, UTX and UTY involve modulation of chromatin accessibility and histone acetylation. This non-enzymatic function likely depends on domains outside the JmjC domain. In UTX, the TPR domain mediates interaction with the MLL complex promoting transcriptional activation (Kato et al., 2020; Rickels et al., 2020), and a corresponding domain in UTY may support similar functions. Additionally, intrinsically disordered regions (IDRs) in UTX contribute to the recruitment of chromatin remodeling factors via liquid–liquid phase separation (Shi et al., 2021). While UTX-IDR drives a more fluid, liquid-like phase, UTY-IDR tends toward a more solid-like behavior. The relatively low expression of UTY in ES cells may serve to prevent this solid-like behavior from interfering with the dynamic chromatin landscape required for maintaining pluripotency. Thus, UTY compensates for its distinct biophysical properties through reduced expression, thereby achieving a functional outcome comparable to UTX. We cannot exclude the possibility that low UTY expression results in sex-based differences in total UTX/UTY dosage, potentially contributing to functional differences between males and females. Indeed, UTX in male ES cells has been linked to the regulation of female-biased gene expression (Ma et al., 2022). Also, we cannot exclude the possibility that residual demethylase activity may contribute to later stages of differentiation, which will require future studies using catalytic-dead models to address directly.

UTX and UTY bind broadly to enhancer regions across the genome; however, only a subset of nearby genes exhibit transcriptional changes upon their loss. Notably, genes such as NODAL and LEFTY1/2, which are rapidly repressed during early differentiation (Besser, 2004), are among those affected, suggesting that UTX and UTY help maintain transcriptional flexibility at select loci. In contrast, many other target genes remain transcriptionally unchanged, with their chromatin structure appearing stable even in the absence of UTX or UTY. This indicates that transcriptional regulation by UTX and UTY is more nuanced than enhancer occupancy alone. Similarly, general coactivators such as p300, BRD4, and the Mediator complex also exhibit widespread binding yet exert locus-specific effects on transcription (Neumayr et al., 2022). These observations suggest that the presence of regulatory complexes at enhancers is not sufficient for gene activation; rather, transcriptional output depends on a permissive chromatin landscape and appropriate molecular partners. This context dependency likely explains why only a subset of UTX/UTY-bound genes show transcriptional responses upon their loss.

Our study also raises important questions about the timing and requirements of UTX and UTY’s catalytic versus non-catalytic functions. In the early stages of development, transcriptional control may rely more on non-enzymatic mechanisms, while enzymatic activity may become essential later for establishing complex transcriptional programs through epigenetic modifications. Thus, functional differences between UTX and UTY may become more pronounced at later stages (Fig. S12). Previous studies have shown UTX’s demethylase roles in cardiac development (Lee et al., 2012) and in the differentiation of skeletal muscle satellite cells (Seenundun et al., 2010) —functions that UTY cannot substitute for. In contrast, UTX functions in a non-catalytic manner during early developmental stages, such as mesodermal differentiation (Wang et al., 2012) and neural progenitor development (Shan et al., 2020), where UTY may compensate. Indeed, our results show that UTX and UTY relocate to the same enhancer regions during early differentiation of ES cells. This suggests that UTY may function in parallel with UTX during the early stages of differentiation. However, as development advances and cellular diversity increases, UTX and UTY may adopt more distinct and cell type–specific roles. It also remains possible that UTY exerts context-dependent enzymatic functions in certain cell types. Clarifying when and how these proteins function—independently or cooperatively—will be key to understanding their roles in regulating gene expression during tissue-specific differentiation.

Recent evidence shows that concurrent loss of UTX and UTY occurs in multiple human cancer types (Gozdecka et al., 2018). This study demonstrates that both proteins modulate enhancer function independently of their enzymatic activity, consistent with our findings in human ES cells. Notably, their combined activity suppresses acute myeloid leukemia, suggesting that UTX and UTY regulate transcriptional programs critical for maintaining ES cell identity and preventing tumorigenesis. Shared mechanisms—such as transcriptional plasticity, chromatin accessibility, and enhancer regulation—imply that UTX and UTY operate through a common regulatory framework in both development and cancer progression.

On the other hand, UTY also exhibits unique regulatory roles in certain tissues, indicating the male-specific regulation. UTY participates in prostate development via chromatin remodeling (Dutta et al., 2016), as well as in regulating gene expression in macrophages and epigenetic changes linked to heart failure progression (Horitani et al., 2024). Notably, forced expression of UTY can masculinize gene expression profiles in the placenta and hypothalamus (Rock et al., 2023). Moreover, UTY influences male-specific DNA methylation in valvular interstitial cells (Gorashi et al., 2025), and promotes spermatogonial proliferation to support meiotic sex chromosome pairing (Subrini et al., 2025). Differences in UTX and UTY expression in brain tissue may underlie sex-biased patterns of gene expression (Xu et al., 2008). Thus, the relative importance of the overlapping and distinct functions of UTX and UTY appears to vary depending on the cell type and developmental context. Such functional versatility may underlie the evolutionary pressure to retain UTY on the Y chromosome.

In summary, our findings demonstrate that UTX and UTY safeguard transcription factor localization, a process essential for maintaining pluripotency. Contrary to the notion that UTY has limited function, our results underscore its broader cooperative role with UTX and highlight the significance of Y-linked genes in regulating transcriptional programs in early development.

## Materials and Methods

### Human ES cell culture and differentiation

The SEES3 hESC line (XY) (Akutsu et al., 2015) was obtained from the Center for Regenerative Medicine, National Research Institute for Child Health and Development (Japan). Cells were cultured feeder-free in StemFit AK02N medium (Takara) on dishes coated with 0.5 μg/cm² Laminin-511 E8 fragment (iMatrix-511; Takara) at 37 °C in a humidified 5% COL atmosphere. For passaging, cells were dissociated using 50% TrypLE Express (Gibco) with 25 mM EDTA in PBS and passaged every 4–5 days before reaching 80% confluency; 10 μM Y-27632 (Wako) was added during passaging to promote survival. H9 (WA09) ES cells (XX) (Thomson et al., 1998) were obtained from the WiCell Research Institute and maintained under the same culture conditions as SEES3 cells. For endoderm differentiation, dissociated cells were seeded in StemFit medium with Y-27632, then switched the next day to STEMdiff Trilineage Endoderm Medium (Veritas) with daily medium changes for three days. Similarly, mesoderm and ectoderm differentiation were performed using the corresponding STEMdiff Trilineage differentiation media. All hESC experiments were approved by the Ethics Committee of Keio University and the Ministry of Education, Culture, Sports, Science and Technology of Japan (October 2012), and by the Ethics Committee of Shiga University of Medical Science for experiments at Yokohama City University (July 2022).

### 3×Flag-HA knock-in at UTX and UTY loci

Human ES cell lines harboring a C-terminal 3×Flag-HA tag at the endogenous UTX or UTY locus were generated using CRISPR-Cas9 genome editing. For UTX, the donor plasmid included 5′ and 3′ homology arms of 952 bp and 825 bp, respectively, flanking the stop codon. For UTY, the 5′ and 3′ homology arms were 1016 bp and 816 bp, respectively. In both cases, a 3×Flag-HA tag followed by a polyadenylation signal was inserted in-frame at the C-terminus within the 3′ UTR, along with a PGK promoter-driven neomycin resistance cassette containing its own polyA signal. The CRISPR-Cas9 was delivered using the pX330-U6-Chimeric_BB-CBh-hSpCas9 plasmid (Addgene #42230, a gift from Feng Zhang) (Cong et al., 2013), which expresses both Cas9 and sgRNA. ES cells were co-transfected with the Cas9-sgRNA and donor plasmids using GeneJuice Transfection Reagent (Merck), following the manufacturer’s instructions. For UTX, the sgRNA target sequence was 5′-AGTACTCTCTCCCGTCCAGT-3′. For UTY, two sgRNAs were used: sgRNA#1, 5′-GGCCATGATGATTACATTTG-3′; and sgRNA#2, 5′-TATTTAATGGCAGTTACGTC-3′. Following selection with 150 µg/ml G418, resistant colonies were isolated and screened by genomic PCR using primers flanking the insertion site. Correct knock-in was confirmed by anti-Flag immunoprecipitation followed by western blotting with anti-UTX or anti-UTY antibodies.

### UTX and UTY knockout cell lines

UTX-KO, UTY-KO, and DKO lines were generated using the CRISPR-Cas9. For each gene, two sgRNAs targeting regions downstream of the ATG start codon were designed and cloned into the pX330 plasmid. For UTX, the sgRNA target sequences were 5′-GCCGCTGCCGCCGCCGCTTT-3′and 5′-GCCGCCGTCACTCGCCCGGT-3′. For UTY, the sgRNA sequences were 5′-ACTACCGCCGCTGTTGCCTT-3′and 5′-CAGACTCCTCTTCACTCTCG-3′. ES cells were co-transfected with the pX330 plasmids and a CAG promoter-driven EGFP-IRES-Puromycin plasmid using GeneJuice. Two days post-transfection, cells were selected with 0.5 µg/ml puromycin and screened for mutations. Genomic DNA was extracted, and the targeted loci were amplified and analyzed by Sanger sequencing to identify insertions or deletions. Clones harboring frameshift mutations resulting in premature stop codons were selected. Two independent clones were established for each genotype. Loss of UTX and UTY protein expression was confirmed by western blotting using anti-UTX and anti-UTY antibodies.

### Dual crosslink ChIP

Cells were cross-linked with 2 mM DSG in PBS/1 mM MgCl_2_ for 40 min at room temperature followed by 1% formaldehyde in PBS for 15 min at room temperature. The reaction was stopped by 125 mM glycine. The cells were washed with PBS and stored at –80°C prior to use. The cells were lysed in Lysis buffer (10 mM Tris-HCl, pH8.0, 1 mM EDTA, 0.5 mM EGTA, 100 mM NaCl, 0.1% Na-Deoxycholate, 0.5% N-lauroyl sarcosine) containing proteinase inhibitor cocktail (Roche). Sonication was conducted with the Handy sonicator UR-21P (Tomy) to generate DNA fragments of approximately 150-450 bp. The sonicated lysates from approximately 2 x 10^6^ cells were diluted in ChIP dilution buffer (10 mM HEPES, pH 7.4, 50 mM NaCl, 1% IGEPAL-CA-630, 10% glycerol) containing proteinase inhibitor cocktail and incubated overnight at 4°C with 50 µl of protein G magnetic beads (Invitrogen) that were preincubated with ∼2 µg of antibodies. The precipitants were washed once with high salt wash buffer (20 mM Tris-HCl, pH8.0, 400 mM NaCl, 2 mM EDTA, 0.1% SDS, 0.2% Triton-X), three times with LiCl wash buffer (10 mM Tris-HCl, pH 8.0, 1 mM EDTA, 250 mM LiCl, 1% NP40, 1% Na-deoxycholate) and once with 10 mM Tris-HCl, pH 8.0, 5 mM EDTA, 10 mM NaCl. Bound chromatin was eluted in elution buffer (90 mM NaHCOL, 1% SDS), and crosslinks were reversed by adding a 1:5 dilution of decross-linking buffer (2 M NaCl, 0.1 M EDTA, 0.4 M Tris-HCl, pH 6.8) and incubating at 65°C for 3 h. Samples were treated with RNase A at 37°C for 30 minutes and proteinase K at 55°C for 1 h. DNA was purified using phenol-chloroform-isoamyl alcohol extraction followed by ethanol precipitation.

### ChIP-seq library preparation and sequencing

ChIP DNA libraries were prepared using the NEBNext ChIP-Seq Library Prep Kit for Illumina (NEB) according to the manufacturer’s instructions. Sequencing was performed on an Illumina HiSeq 2500 platform (50 bp, single-end).

### Processing and analysis of ChIP-seq data

Raw sequencing reads were quality-controlled and trimmed using Fastp to remove adapter sequences and low-quality bases. The resulting reads were aligned to the human reference genome (GRCh38) using Bowtie2 with default parameters. Aligned reads were processed using SAMtools to generate BAM and index files. Peak calling was performed with MACS2 in narrowPeak mode using a q-value cutoff of < 0.05. For each condition, peaks identified from three independent biological replicates were merged, retaining only those located on autosomes (chromosomes 1–22) and sex chromosomes (X and Y). Peaks detected uniquely in only one replicate with abnormally high signal were excluded. BAM files from the three replicates were merged for downstream analysis. Differential peak analysis was conducted using MAnorm with default parameters. To compare peak locations between conditions, genomic distances were computed using bedtools closest. For visualization, bamCoverage was used to generate BigWig files with counts per million normalization. These tracks were visualized using the Integrative Genomics Viewer. Signal differences between conditions were assessed using bigwigCompare. Heatmaps and aggregation plots were created using deepTools computeMatrix, followed by visualization with plotHeatmap and plotProfile. De novo and known motif enrichment analyses were performed using HOMER with a 200 bp window size and the -mask option to exclude repetitive sequences. Genomic annotations of peaks relative to TSS and gene bodies were obtained using HOMER annotatePeaks.pl. Gene set enrichment analysis (GSEA) was conducted to assess the association between OCT4 binding and gene expression changes.

### RNA-seq library preparation and sequencing

RNA-seq libraries were prepared from 500 ng of total RNA per sample using the NEBNext Poly(A) mRNA Magnetic Isolation Module, followed by the NEBNext Ultra II Directional RNA Library Prep Kit for Illumina (NEB), according to the manufacturer’s protocols. Sequencing was performed on an Illumina HiSeq 2500 platform using single-end 50 bp reads.

### Processing and analysis of RNA-seq data

Sequence reads were aligned to the human genome (GRCh38) using STAR. TPM and FPKM values were estimated using RSEM. Gene-level expression was also quantified using featureCounts based on the annotation file Homo_sapiens.GRCh38.105.gtf. For differential expression analysis, raw counts obtained by featureCounts were analyzed using the TCC-GUI pipeline in R. Two independent biological replicates were used for comparisons. Normalization was performed using the TMM method, and differentially expressed genes were identified using edgeR. Genes with a logL fold change > 1 and a p-value < 0.05 were considered significantly differentially expressed. Gene ontology enrichment analysis was performed using the ToppGene Suite (https://toppgene.cchmc.org/). Principal component analysis was performed using ExAtlas (https://kolab.elixirgensci.com/exatlas/).

### ATAC-seq

ATAC-seq was performed using the Omni-ATAC protocol (Corces et al., 2017) with minor modifications. Briefly, 1×10L viable cells were resuspended in 50 µl of cold ATAC Resuspension Buffer (RSB)(10 mM Tris-HCl (pH 7.4), 10 mM NaCl, and 3 mM MgClL) containing 0.1% NP-40, 0.1% Tween-20, and 0.01% digitonin. After incubation on ice for 3 min, cells were washed with 1 ml of cold RSB containing 0.1% Tween-20, and nuclei were pelleted at 500 × g for 10 min at 4°C. The nuclear pellet was resuspended in 50 µl of transposition mix (25 µl of 2× TD buffer (20 mM Tris-HCl, pH 7.6; 10LmM MgClL; 20% dimethylformamide), 2.5 µl of Tn5 transposase, 16.9 µl of PBS, 0.5 µl of 10% Tween-20, 0.1Lµl of digitonin, and 5 µl of water), and incubated at 37°C for 30 min in a thermomixer at 630 rpm. Transposed DNA was purified with AMPure XP beads (1.8×) and eluted in 20 µl of EB buffer. Libraries were pre-amplified with NEBNext High-Fidelity 2X PCR Master Mix for 5 cycles, and 10% of the reaction was used in a qPCR side reaction to determine the required number of additional cycles. Final amplification was performed with the remaining reaction, and PCR products were purified using a two-step AMPure XP bead cleanup (0.6× followed by 1.3×). Final libraries were eluted in 20 µl of EB buffer and assessed using a Bioanalyzer (Agilent) to confirm fragment size distribution.

### Teratoma formation assay

Human ES cells were dissociated into single cells and resuspended in a 1:2 mixture of Matrigel (Corning) and StemFit AK02N medium supplemented with 10 µM Y-27632. A total of 1 × 10L cells in 100 µL of this mixture were subcutaneously injected into the dorsal flank of immunodeficient SCID mice (CLEA Japan). After 8 weeks, the mice were sacrificed, and the injection sites were dissected to assess teratoma formation. The excised tissues were fixed in 4% paraformaldehyde, embedded in paraffin, and sectioned for histological and immunohistochemical analyses. All animal procedures were approved by the Animal Care and Use Committee of Keio University School of Medicine.

### Quantification and statistical analysis

Descriptive and comparative statistics were performed as detailed in the figure legends with the number of replicates indicated. Significance is defined as a p-value less than 0.05 indicated with asterisk. Error bar represents the standard deviation of the mean of the replicates. Statistical analyses were implemented with Prism 9 (GraphPad) or Excel (Microsoft) software.

## Supporting information

Supplemental_Figures_and_Table_revised

## Data availability

All sequencing data have been deposited in the GEO database under series accession number GSE301298.

## Oligonucleotides and antibodies

The oligonucleotide primers and antibodies used in this study are listed in Supplemental Table S1.

## Acknowledgments

We thank the members of our laboratory and the Collaborative Research Resources, Keio University, for technical assistance. We also thank Hitoshi Niwa of Kumamoto university for kindly providing the SOX2 antibody. This work was supported by grants from the Japan Society for the Promotion of Science (JSPS) Grant-in-Aid for Specially Promoted Research (JP19K06492, JP20H05395, JP20H04929, JP22K06090, JP22H04699 to T.A.).

## Author contributions

T.A. conceived the project, performed the experiments, and analyzed the data. S.S. and M.Y. performed experiments, and T.N. and M.Y. analyzed the NGS data. K.I. contributed to the generation of knock-in cells. H.T. contributed to the data analysis. T.A. and M.S.H.K. supervised the project and interpreted the results. T.A. wrote the manuscript with feedback from co-authors.

