## Supplemental_Figures_and_Table_revised for "Functional redundancy between UTY and UTX in regulating the localization of transcription factors involved in pluripotency"

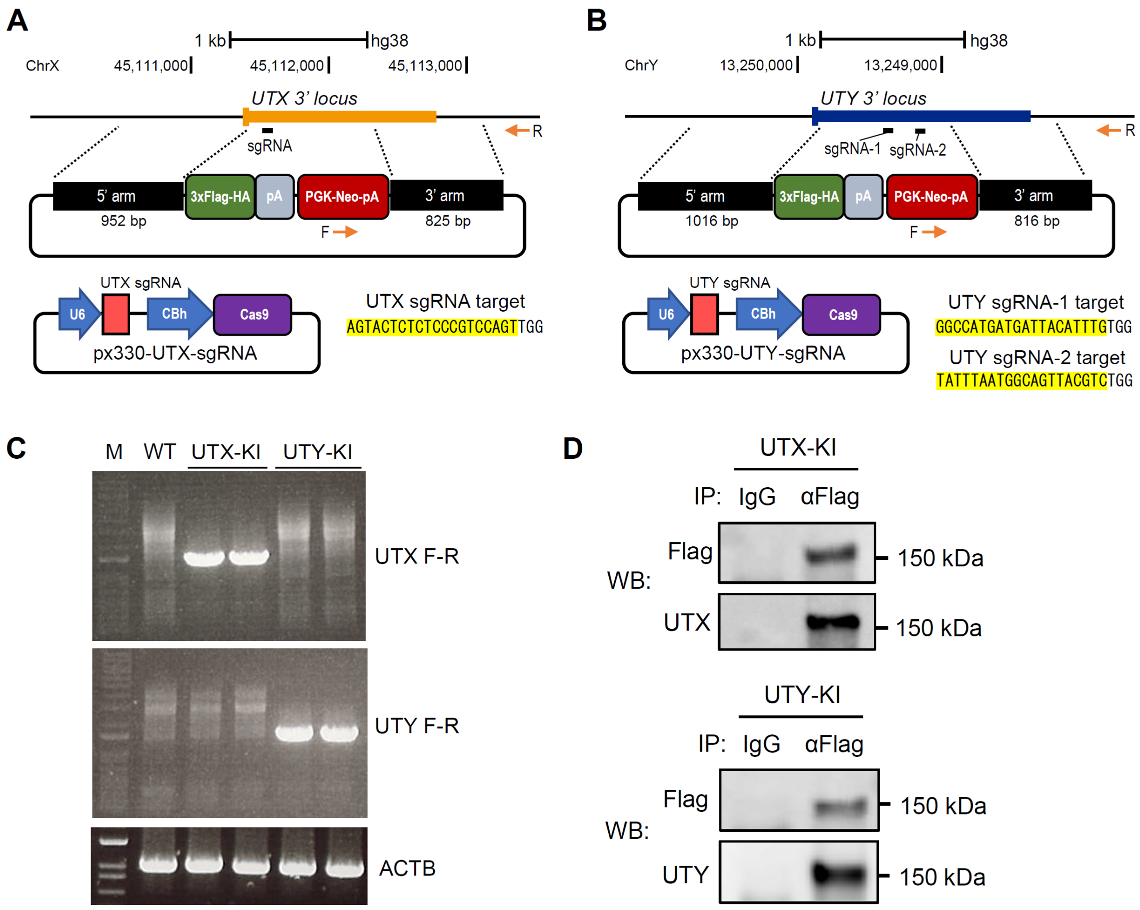


**Fig. S1. Generation of UTX-KI and UTY-KI human ES cell lines**

(A) Schematic diagram of UTX-KI design. The 3xFlag-HA tag and poly A signal were inserted at the C-terminus of UTX. 5' and 3' homology arm sequences were designed for homologous recombination (HR). A PGK promoter-driven Neomycin resistance gene cassette with a poly A signal was also included for drug selection. CRISPR-Cas9-mediated gene editing was performed using the px330 vector, and the target sequences for the sgRNAs are indicated.

(B) Schematic diagram of UTY-KI design. Similar to UTX-KI cells, the UTY gene was modified by introducing a 3xFlag-HA tag and poly A signal at the C-terminus. Two target sites for the sgRNAs were designed for the generation of UTY-KI cells.

(C) Genomic PCR confirming successful knock-in at each locus. PCR was performed with a pair of forward primer (F) and reverse primer (R) specific to each locus, as shown in panels (A) and (B), respectively. ACTB was used as a control for the PCR amplification.

(D) Flag immunoprecipitation followed by western blot using anti-UTX or anti-UTY antibodies, confirming that the tagged protein corresponds to the target protein.

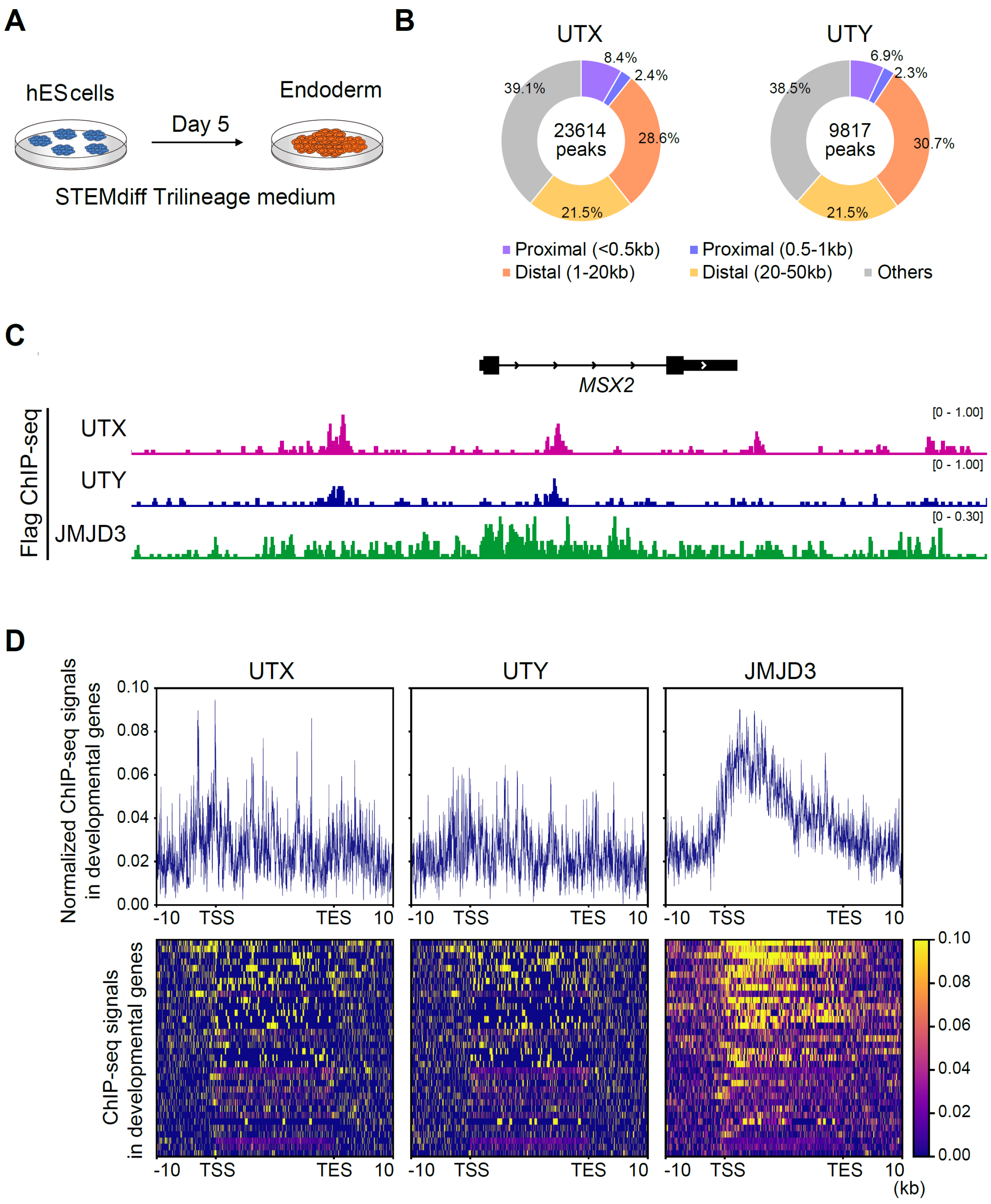


**Fig. S2. Localization patterns of UTX, UTY, and JMJD3 in differentiated cells**

(A) Human ES cells were differentiated into endoderm cells by culturing in STEMdiff Trilineage medium for 5 days.

(B) Number of UTX and UTY ChIP-seq peaks and their distribution relative to the transcription start site (TSS).

(C) Flag ChIP-seq showing the distribution of UTX, UTY, and JMJD3 at the *MSX2* locus, a lineage-associated developmental gene.

(D) Flag ChIP-seq showing the distribution of UTX and UTY at developmental gene loci, distinct from JMJD3, which is localized from TSS to TES.

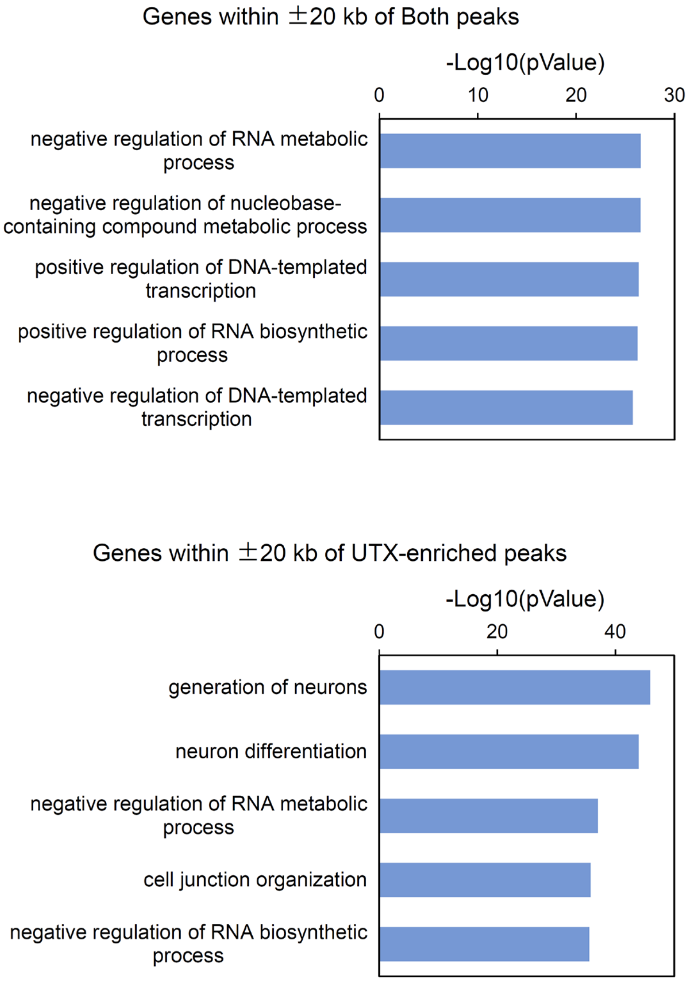


**Fig. S3. Gene ontology analysis of genes associated with UTX/UTY co-bound and UTX-enriched binding sites**

(A) Genes located within ±20 kb of UTX/UTY co-bound sites (Both sites).

(B) Genes located within ±20 kb of UTX-enriched sites.

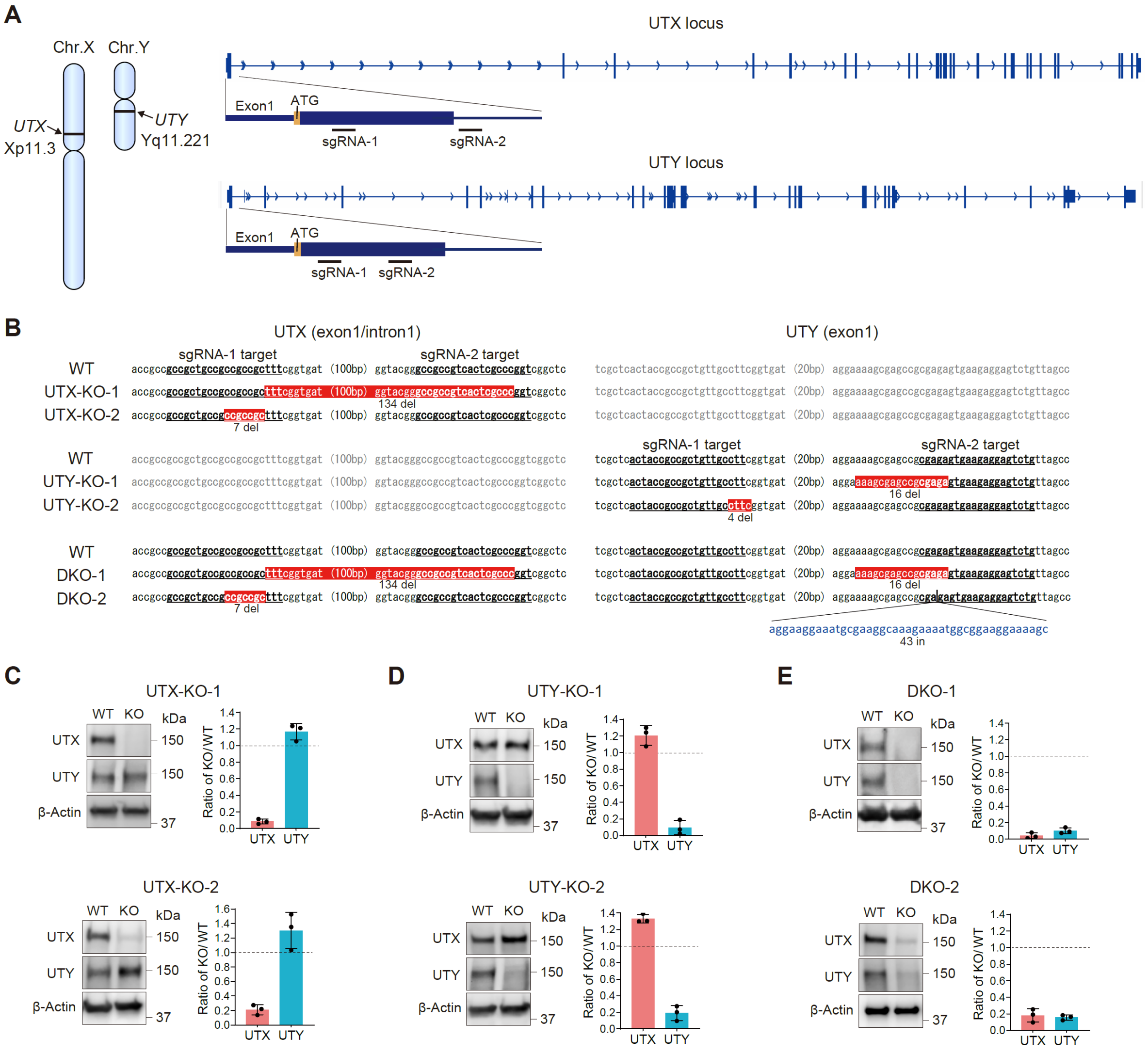


**Fig. S4. Generation of UTX-KO, UTY-KO, and DKO human ES cell lines**

(A) Locations of the UTX and UTY genes on the XY chromosomes. CRISPR-sgRNAs were designed at two target sites for each gene to generate knockouts.

(B) Mutations in UTX-KO, UTY-KO, and DKO cell lines induced by CRISPR-Cas9 editing. Two clones were generated for each knockout. "del" indicates deletions, and "in" indicates insertions.

(C–E) Western blot analysis confirming the absence of the target proteins in UTX-KO (C), UTY-KO (D), and UTX/UTY double-knockout (DKO) (E) cell lines. Protein signals were normalized to β-actin, and KO/WT ratios were calculated based on the normalized band intensities.

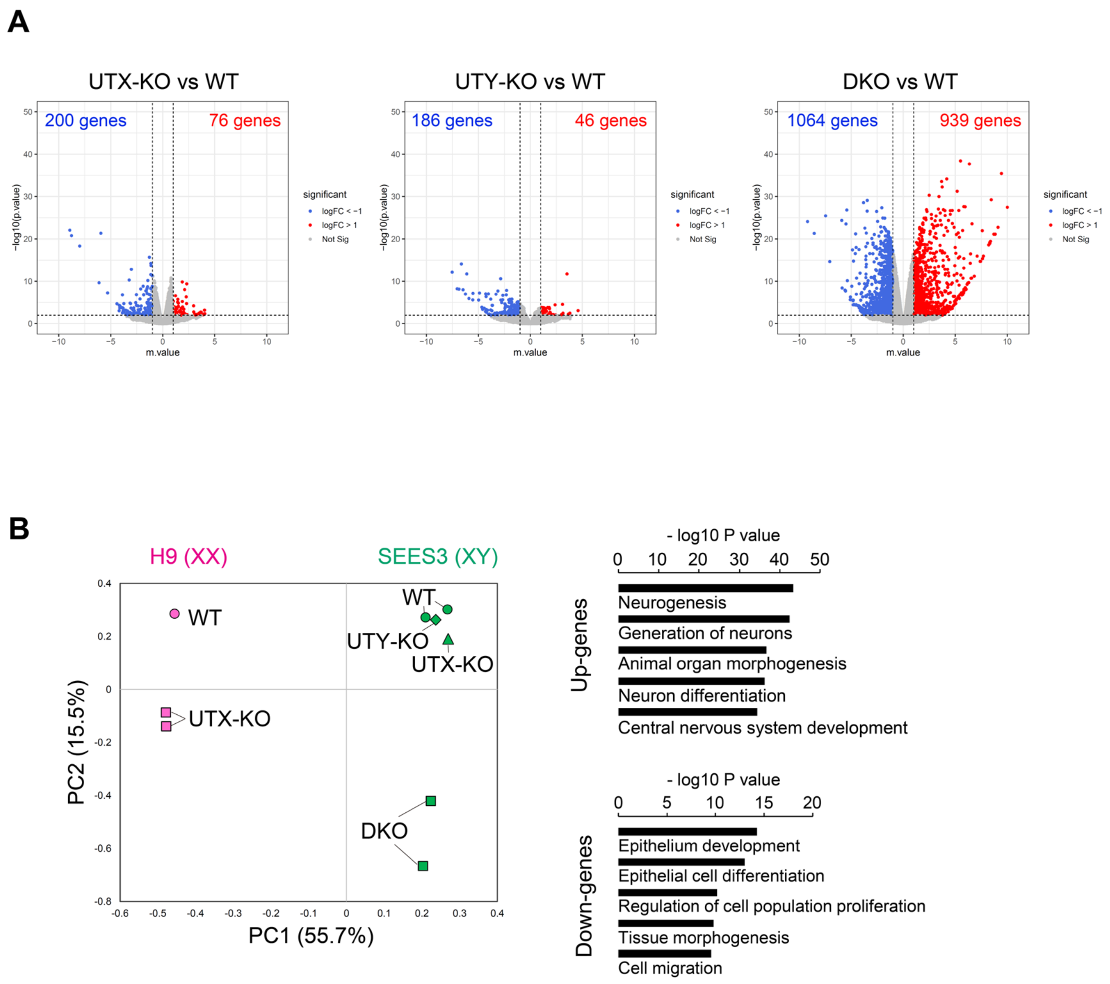


**Fig. S5. Gene expression changes induced by UTX-KO, UTY-KO, and DKO and comparison of male (XY) DKO with female (XX) UTX-KO ES cells**

(A) Volcano plots showing differential gene expression in UTX-KO, UTY-KO, and DKO cells based on transcriptome-wide mRNA-seq analysis of all genes. Blue and red dots indicate downregulated and upregulated genes, respectively.

(B) Principal component analysis of male (XY) WT, UTX-KO, UTY-KO, and DKO, and female (XX) WT and UTX-KO ES cells. PC1 largely represents Y chromosome–linked gene expression, while PC2 shows a similar shift in male DKO and female UTX-KO cells.

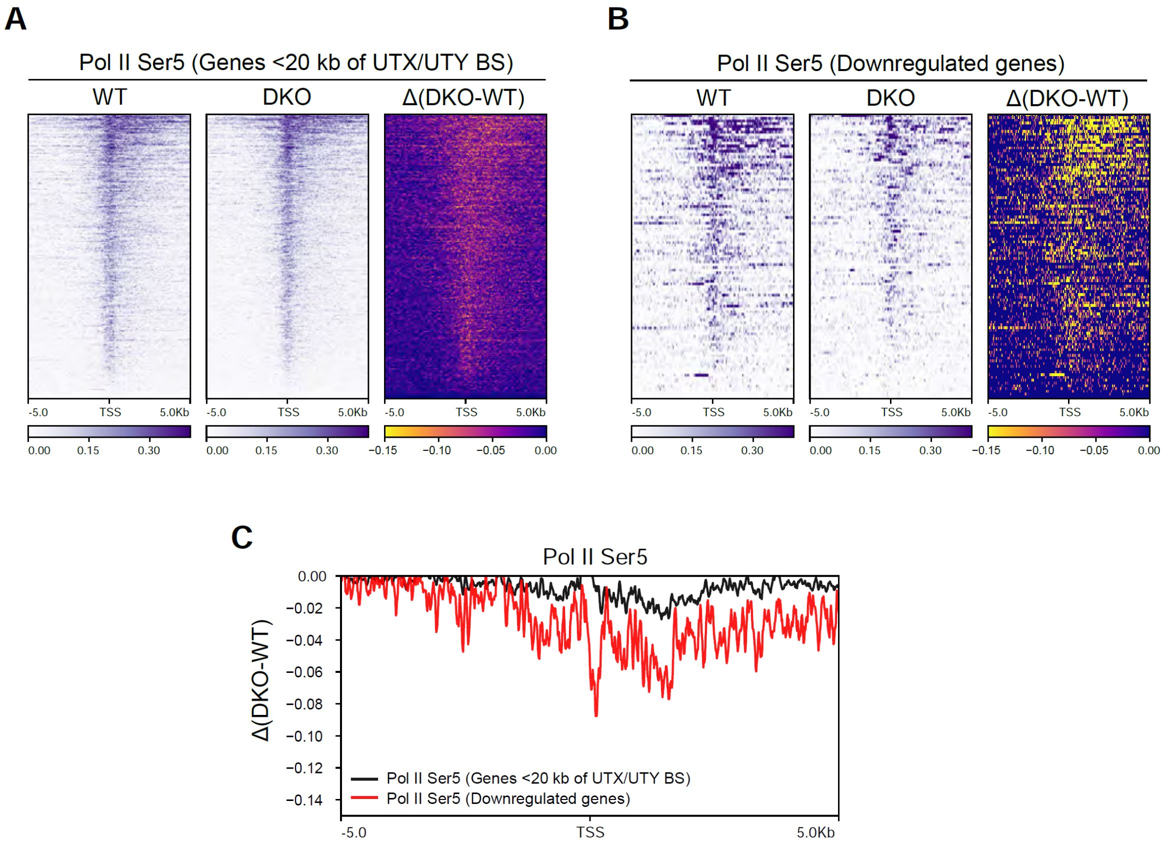


**Fig. S6. UTX/UTY DKO results in a reduction of RNA polymerase II (Pol II) loading**

(A) Pol II Ser5 ChIP-seq signals in WT and DKO cells within 10 kb of the TSS of genes associated with UTX/UTY binding sites. Right panel shows differences in the signals between WT and DKO.

(B) Pol II Ser5 ChIP-seq signals in WT and DKO cells within 10 kb of the TSS of genes associated with UTX/UTY binding sites and whose expression is suppressed in DKO cells. The right panel shows the differences in signal intensity between WT and DKO.

(C) The reduction in phosphorylation at Ser5 is greater in genes that are downregulated in DKO cells.

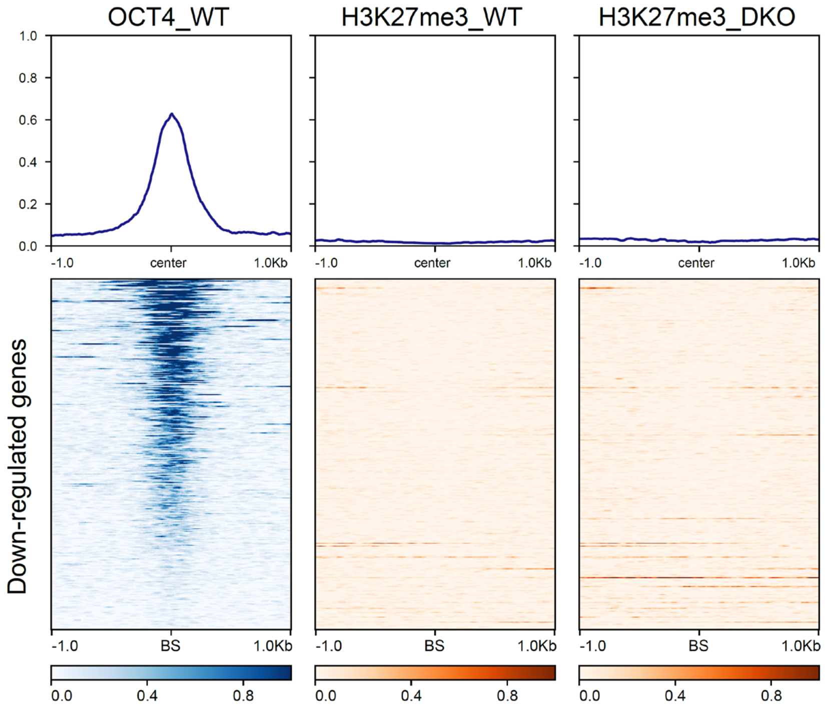


**Fig. S7. OCT4 and H3K27me3 profiles in WT and DKO cells at DKO-upregulated genes**
OCT4 and H3K27me3 ChIP-seq signal profiles in WT and DKO cells centered on UTX/UTY binding sites near genes downregulated in DKO cells (same sites as in Figure 4E).

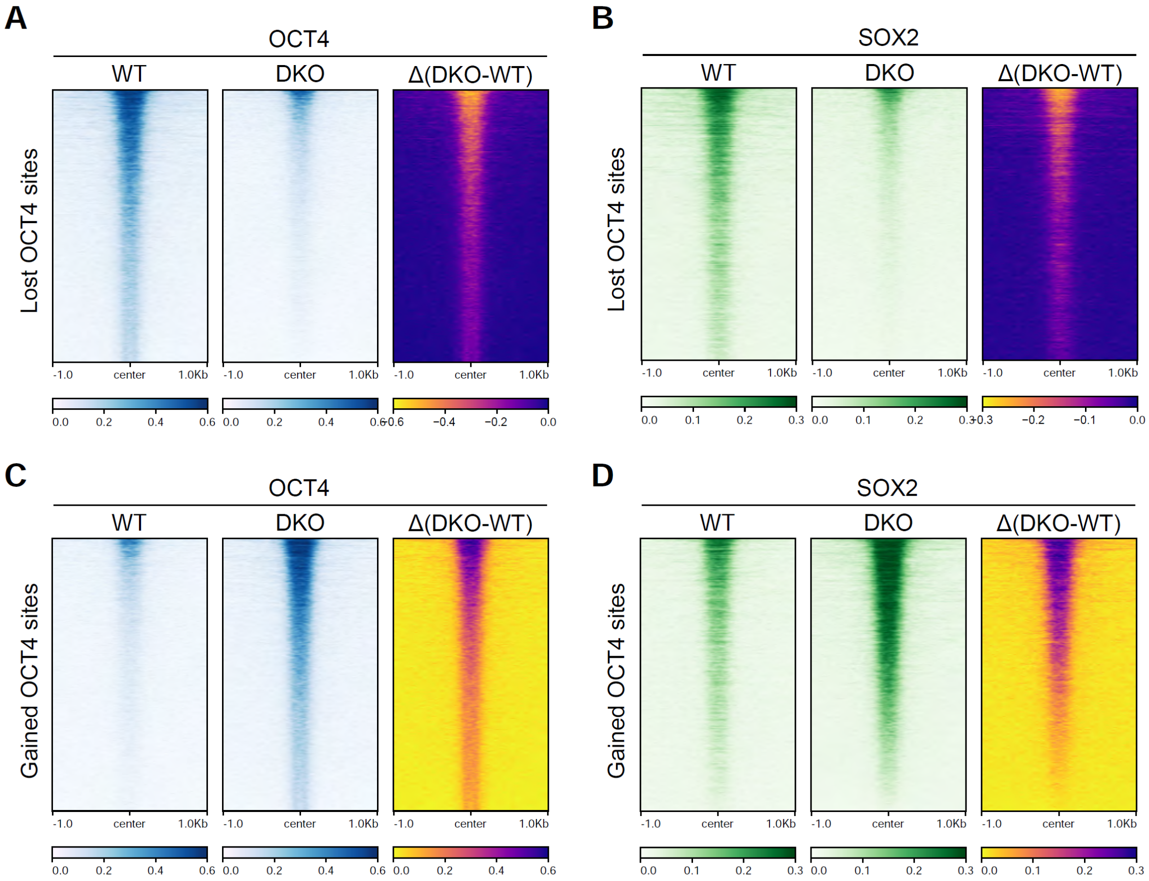


**Fig. S8. UTX/UTY DKO results in the relocation of SOX2 as well as OCT4**

(A) ChIP-seq signal intensities of OCT4 at OCT4-lost sites.

(B) Comparison of SOX2 ChIP-seq signal intensities between WT and DKO cells within a 2 kb region centered on OCT4-lost sites.

(C) ChIP-seq signal intensities of OCT4 at OCT4-gained sites.

(D) Comparison of SOX2 ChIP-seq signal intensities between WT and DKO cells within a 2 kb region centered on OCT4-gained sites.

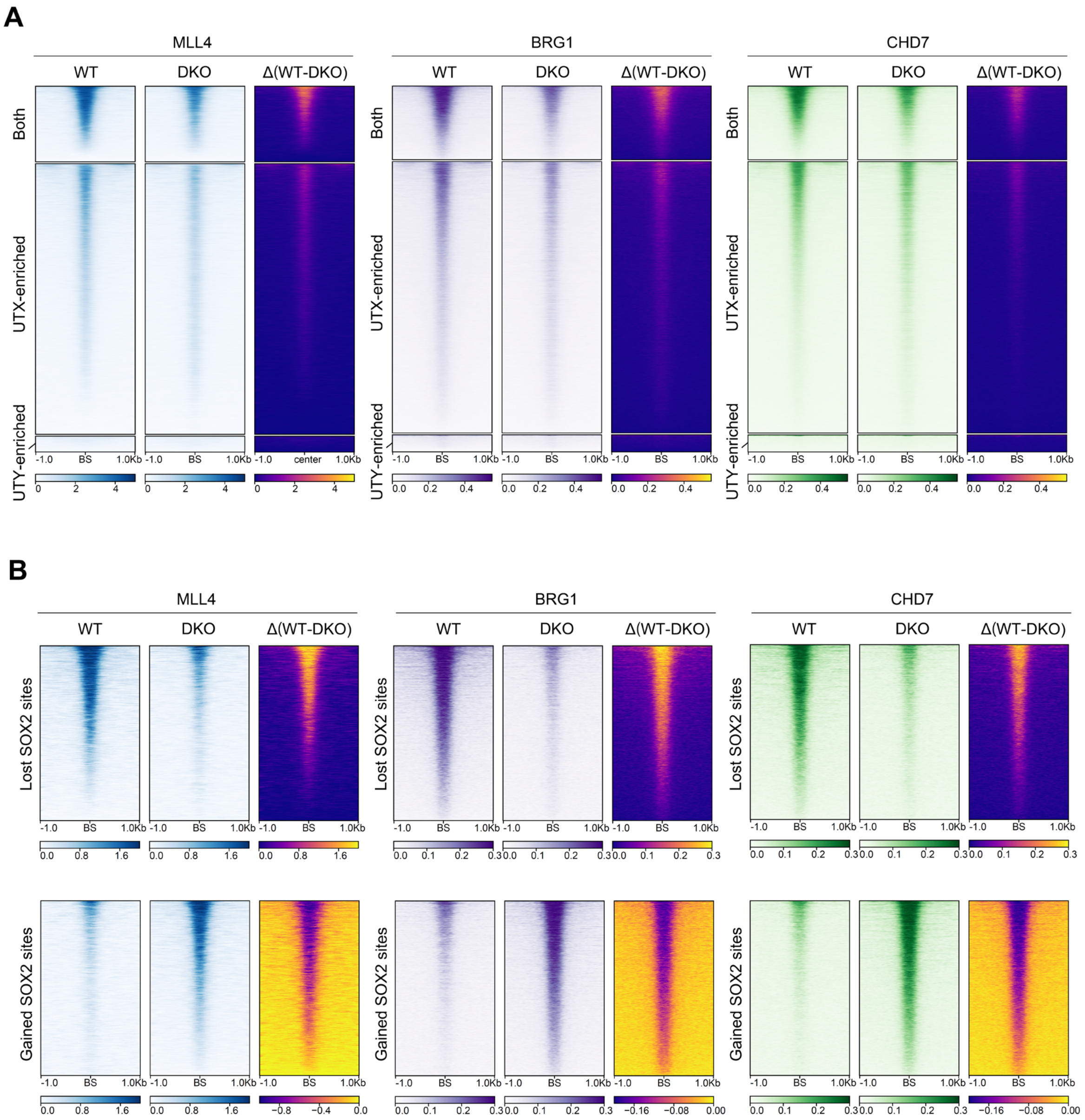


**Fig. S9. UTX and UTY are required for the distribution of MLL4, BRG1, and CHD7**

(A) Comparison of ChIP-seq signal intensities of the indicated factors between WT and DKO cells centered on Both BS (UTX/UTY co-bound sites), UTX-enriched BS, and UTY-enriched BS.

(B) Comparison of ChIP-seq signal intensities of the indicated factors between WT and DKO cells centered on lost and gained SOX2 binding sites.

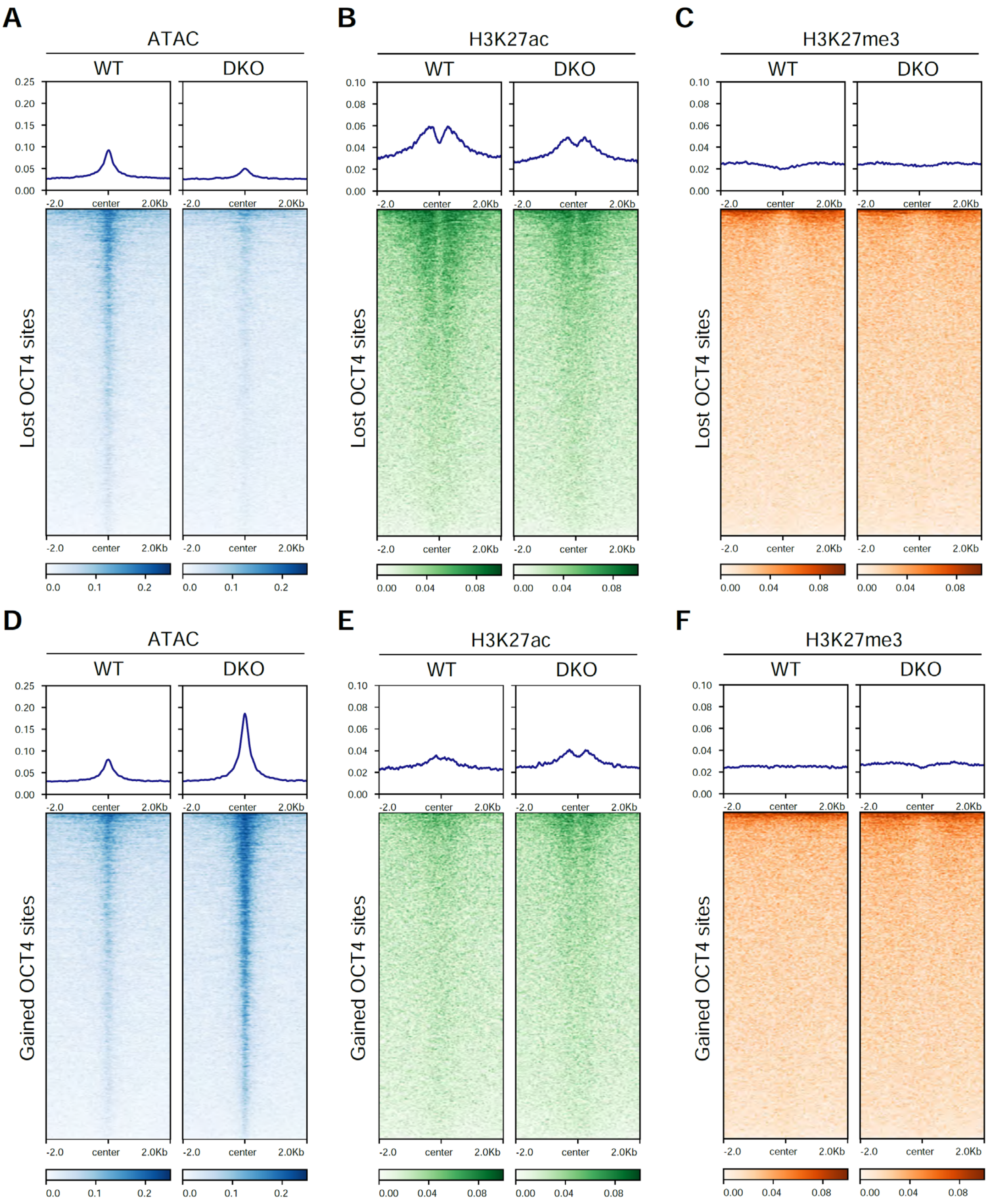


**Fig. S10. Changes in chromatin accessibility and histone modifications following UTX/UTY DKO**

(A-C) Comparison of ATAC-seq, H3K27ac ChIP-seq, and H3K27me3 ChIP-seq signal intensities between WT and DKO cells within a 2 kb region centered on OCT4-lost sites.

(D-F) Comparison of ATAC-seq, H3K27ac ChIP-seq, and H3K27me3 ChIP-seq signal intensities between WT and DKO cells within a 2 kb region centered on OCT4-gained sites.

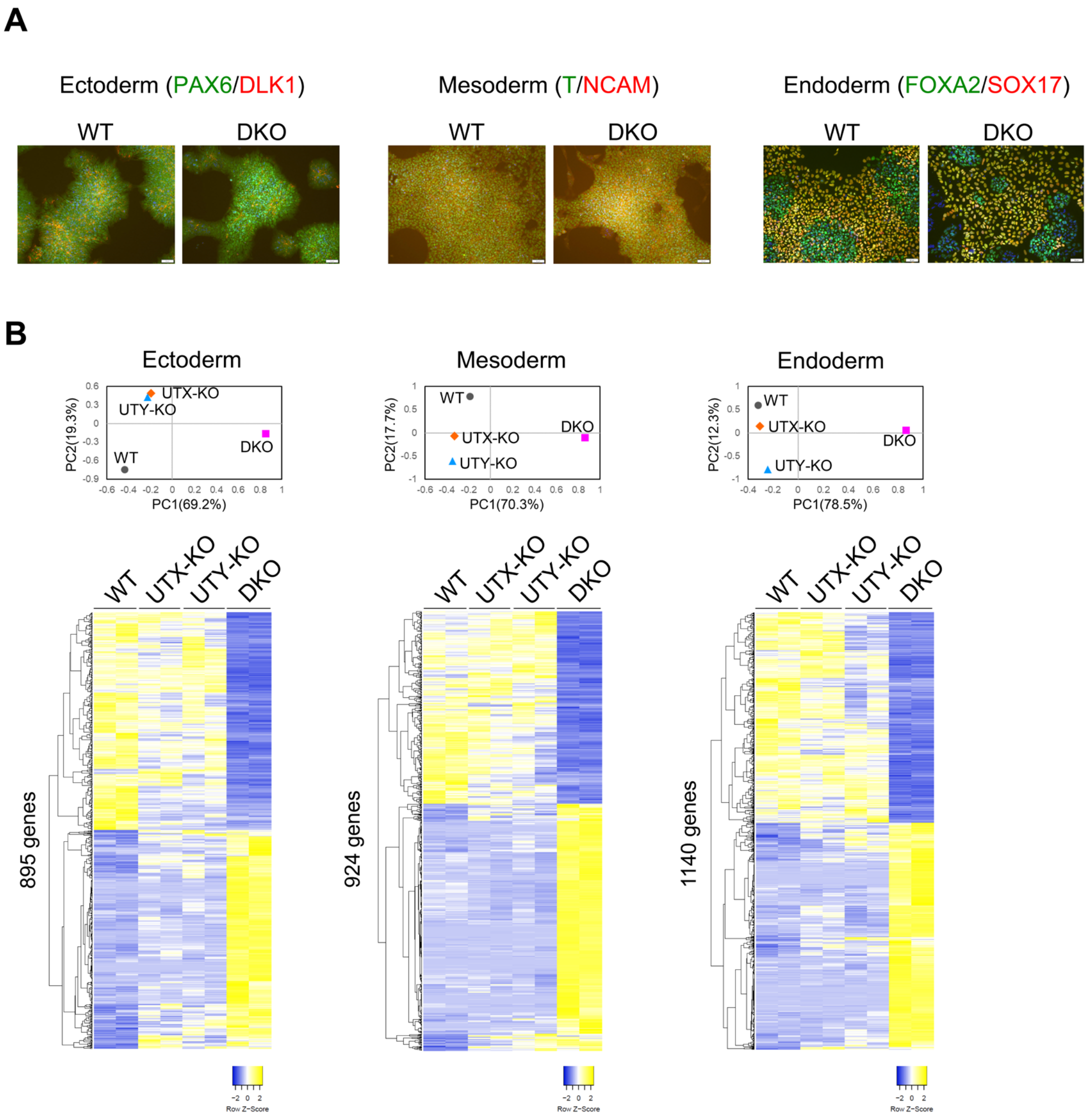


**Fig. S11. Effects of UTX and UTY loss on three-germ-layer differentiation in vitro**

(A) Immunostaining analysis of representative markers for the three germ layers in differentiated cells derived from WT and DKO ES cells.

(B) PCA and heatmaps showing differential gene expression in the differentiated cells derived from WT, UTX-KO, UTY-KO, and DKO ES cells.

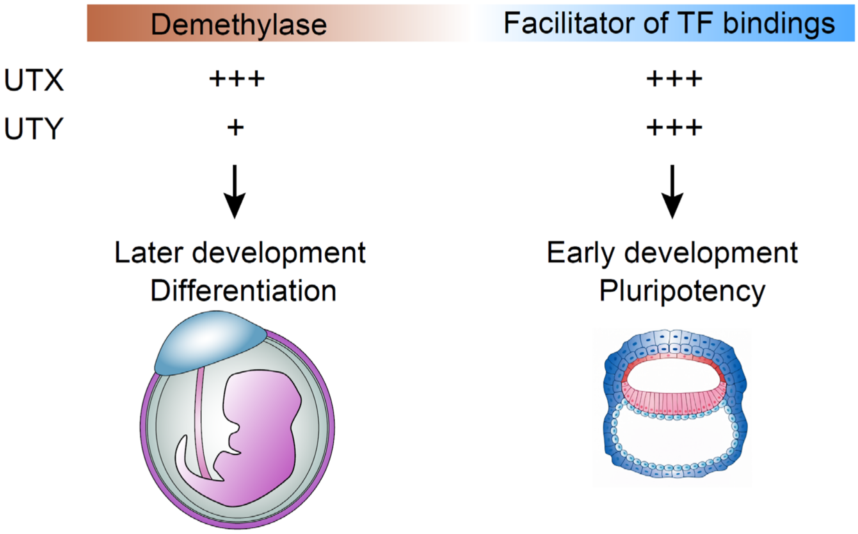


**Fig. S12. Functional divergence and overlap between UTX and UTY during development**

UTX and UTY possess two functions: H3K27 demethylase activity and facilitation of transcription factor (TF) binding. While they exhibit a clear difference in demethylase activity, they have a comparable ability to promote TF binding (a finding in this paper). Demethylase activity primarily contributes to later developmental stages and differentiation, whereas facilitation of TF binding is more relevant during early development and pluripotency maintenance. UTY compensates for the loss of UTX by maintaining enhancer accessibility and transcriptional programs critical for early development.

### Supplemental Table S1. Oligonucleotides and antibodies

**KI-arm Primers**

| Name | Forward Sequence | Reverse Sequence |
|---|---|---|
| KI-UTX-5arm | CTGGGTGAGACCAGCCTAAA | AGATGAGGCGGATGGTAATG |
| KI-UTX-3arm | CAGTGCCCAGTTTTTGTTCA | AATCAGCAGGGCATTTCACT |
| KI-UTY-5arm | AGCCATCCTCGTATTTCACG | AGATGAGGATGATAATGAAA |
| KI-UTY-3arm | AAGGGTTTCCCAATGTTTGA | CAATGATCAAGACATTTCACC |
| KI-JMJD3-5arm | GCCCCGAGTGTTACTGTGTT | TCGCGACGTGCTGGCTGGGG |
| KI-JMJD3-3arm | ATGCTAGCTGTGACCCCTGT | CCCGGTACTCAACCTGCTAC |

**guideRNA Oligos**

| Name | Forward Sequence | Reverse Sequence |
|---|---|---|
| UTX-KI-gRNA | caccGAGTACTCTCTCCCGTCCAGT | aaacACTGGACGGGAGAGAGTACTC |
| UTY-KI-gRNA-1 | caccGGCCATGATGATTACATTTG | aaacCAAATGTAATCATCATGGCC |
| UTY-KI-gRNA-2 | caccGTATTTAATGGCAGTTACGTC | aaacGACGTAACTGCCATTAAATAC |
| JMJD3-KI-gRNA | caccGGCCCCGCGGCGCCTTGCGC | aaacGCGCAAGGCGCCGCGGGGCC |
| UTX-KO-gRNA-1 | caccGCCGCTGCCGCCGCCGCTTT | aaacAAAGCGGCGGCGGCAGCGGC |
| UTX-KO-gRNA-2 | caccGCGCCGCCGCTTTCGGTGATG | aaacCATCACCGAAAGCGGCGGCGC |
| UTY-KO-gRNA-1 | caccGACTACCGCCGCTGTTGCCTT | aaacAAGGCAACAGCGGCGGTAGTC |
| UTY-KO-gRNA-2 | caacGCAGACTCCTCTTCACTCTCG | aaacCGAGAGTGAAGAGGAGTCTGC |

**RT-qPCR Primers**

| Gene | Forward Sequence | Reverse Sequence |
|---|---|---|
| CDH17 | TCCTCGGACCAGATCGAAATG | GCGGGACTTAACGTCCAGT |
| LGALS4 | CCCTTCTATGAGTACGGGCAC | TGAAGTTGCAGATCCCCATCC |
| MUC13 | CAGACAGTGAGTCAACCACAAA | GGACCTGTGCTGTTTAGGGT |
| HMGCS2 | GCCCAATATGTGGACCAAACT | GAAGCCCATACGGGTCTGG |
| TM4SF4 | CTGTGGTTGGATTCTTGGGAG | GGGTAGCCCCATGTACTATTGG |
| GLTPD2 | TCAATTCCTCTCGCTATCTTCGC | CACATCCCCTTCGGGTTTCA |
| CLDN18 | ACATGCTGGTGACTAACTTCTG | AAATGTGTACCTGGTCTGAACAG |
| PAX2 | CACTTGCGAGCTGACACCTT | TGCAGATAGACTCGACTTGACTT |
| TMEM72 | AGGGGCCTACTTTGTGGCT | TTCTCCCTTACTCTGTCTGCC |
| MYOG | GGGGAAAACTACCTGCCTGTC | AGGCGCTCGATGTACTGGAT |
| TNNT3 | AGGAGCTGGTCGCTCTCAA | CCTTCTCTGCACGAATCCTCT |
| SGCD | AATCCAGTCCCGACCAGGTAA | TCCGCTCCTAAAACTCGTAATCT |
| LSP1 | GGAGCACCAGAAATGTCAGCA | TCGGTCCTGTCGATGAGTTTG |
| MAP2 | CCAATGGATTCCCATACAGG | TCTCCGTTGATCCCATTCTC |
| GAP43 | GGCCGCAACCAAAATTCAGG | CGGCAGTAGTGGTGCCTTC |
| CBLN2 | AACCGCACCATGACCATCTAT | GCAAGATCAAAGTGGTTGCCAAT |
| SLC39A2 | TGGACATTTACCCTCACCCTC | CACAAGCCCCTTATGAGCCA |
| KRT15 | TCTGCTAGGTTTGTCTCTTCAGG | CCAGGGCACGTACCTTGTC |

| Antibody | Source | Identifier | Application(s) |
|---|---|---|---|
| UTX | Cell Signaling | 33510 | WB |
| UTY | Abcam | ab91236 | WB |
| Flag | Sigma | F1804 | ChIP, ICC |
| HA | Abcam | ab9110 | WB |
| ACTB | Cell Signaling | 4970 | WB |
| PolII S5 | Abcam | AB5408 | ChIP |
| OCT4 | Abcam | AB19857 | ICC, ChIP, WB |
| SOX2 | Provided by Hitoshi Niwa | | ChIP |
| H3K27ac | Active Motif | 39133 | ChIP |
| H3K27me3 | Cell Signaling | 07-449 | ChIP |
| H3 | Abcam | ab1791 | WB |
| MLL4 | Merk | ABE1867 | ChIP |
| BRG1 | Abcam | Ab110641 | ChIP |
| CHD7 | Cell Signaling | 6505S | ChIP |
| CDH17 | Sigma | HPA026556 | IHC |
| ACTN2 | Abcam | ab68167 | IHC |
| TUBB3 | Abcam | ab18207 | IHC |
| NESTIN | R&D | MAB1259 | ICC |
| PAX6 | Abcam | ab195045 | ICC |
| DLK1 | Abcam | ab89908 | ICC |
| T | R&D | AF2085 | ICC |
| NCAM | Santa Cruz | sc-7326 | ICC |
| FOXA2 | R&D | AF2400 | ICC |
| SOX17 | Cell Signaling | 81778S | ICC |

WB: Western Blot
ChIP: Chromatin Immunoprecipitation
ICC: Immunocytochemistry
IHC: Immunohistochemistry
